# Intrinsic dynamics enhance temporal stability of stimulus representation along rodent visual cortical hierarchies

**DOI:** 10.1101/822130

**Authors:** Eugenio Piasini, Liviu Soltuzu, Paolo Muratore, Riccardo Caramellino, Kasper Vinken, Hans Op de Beeck, Vijay Balasubramanian, Davide Zoccolan

## Abstract

Along the ventral stream, cortical representations of brief, static stimuli become gradually more invariant to identity-preserving transformations. In the presence of long, temporally structured dynamic stimuli, higher invariance should imply temporally persistent representations at the top of this functional hierarchy. However, such stimuli could engage adaptive and predictive processes, whose impact on neural coding dynamics is unknown. By probing the rat analogue of the ventral stream with movies, we uncovered a hierarchy of temporal scales, with deeper areas encoding visual information more persistently. Furthermore, the impact of intrinsic dynamics on the stability of stimulus representations gradually grew along the hierarchy. Analysis of a large dataset of recordings from the mouse visual hierarchy yielded similar trends, revealing also their dependence on the behavioral state of the animal. Overall, these findings show that visual representations become progressively more stable along rodent visual processing hierarchies, with an important contribution provided by intrinsic processing.

## Introduction

The visual system of primates and other mammals is able to support discrimination and categorization of visual objects despite major variation in their appearance, resulting (for instance) from translations, rotations and changes of scale [1–3]. Decades of research on non-human primates have shown that such ability (referred to as *invariant object recognition*) is the result of processing along the ventral visual stream – a hierarchy of visual cortical areas, along which invariance to object transformations emerges progressively as the outcome of largely feedforward computations [1; 4; 5]. While neurons at early stages of this pathway (e.g., simple cells in primary visual cortex; V1) are highly sensitive to variation in the appearance of their preferred visual stimuli (e.g., the position of oriented edges), units at the apex of the processing hierarchy (the anterior inferotemporal cortex; IT) maintain their tuning for visual objects despite (e.g.) position and scale changes. This allows visual IT neurons to better support invariant object recognition, as compared to lower-level visual areas [6–10]. However, these conclusions mainly derive from studies relying on presentation of brief, static visual stimuli, appositely designed to engage feedforward processing, while minimizing the influence of recurrent and top-down processing, as well as the impact of adaptation. As a result, it remains unknown how the build-up of invariance along the ventral stream would emerge under more naturalistic viewing conditions, such as in the presence of dynamic visual stimuli (e.g., natural movies lasting at least several seconds), when an uninterrupted flow of visual information hits the retina.

Under such naturalistic stimulation, two alternative scenarios are conceivable. One possibility is that the increase of invariance along the ventral stream (as tested with static objects) may directly translate into a concomitant increase of temporal stability of neuronal responses (as tested with natural movies). This equivalence between invariance and temporal stability can be easily understood by thinking about an idealized invariant object detector (Fig. 1). Such idealized unit would respond to the presence of its preferred object (the “rat head” in this example) within its receptive field (RF; yellow shape in Fig. 1a) regardless of the specific position, size or orientation of the object. As a result, if the object happened to enter the unit’s RF while viewing a dynamic scene, the unit would start firing to report its presence and would keep firing at approximately the same rate as long as the object remained inside the RF (green trace in Fig. 1b, top), while smoothly transforming (e.g., translating and/or rotating) along the way. By contrast, a poorly invariant, lower-level unit (e.g., a V1-like edge detector) would fire more transiently (green trace in Fig. 1b, bottom), given that features matching its selectivity would quickly enter and exit its much smaller RF (orange circle in Fig. 1a), while the movie unfolds over time. The tight relationship between invariance and temporal stability (or slowness) is exploited in many theoretical accounts of how invariance is achieved along the ventral stream, where temporal continuity of the visual input is the key statistical structure exploited by unsupervised learning mechanisms to build invariant representations [11–19]. These unsupervised temporal learning processes are typically instantiated in purely feedforward architectures. More in general, the equivalence between invariance and temporal stability is consistent with any feedforward processing hierarchy (such as standard convolutional neural networks [20]), in which the instantaneous response of each unit to the current visual input only depends on the input itself and is not affected by the previous activity of the unit or the current state of the network. In this scenario, an object smoothly transforming within a dynamic scene is treated and processed as a sequence of static, independent snapshots.

**Fig. 1.**
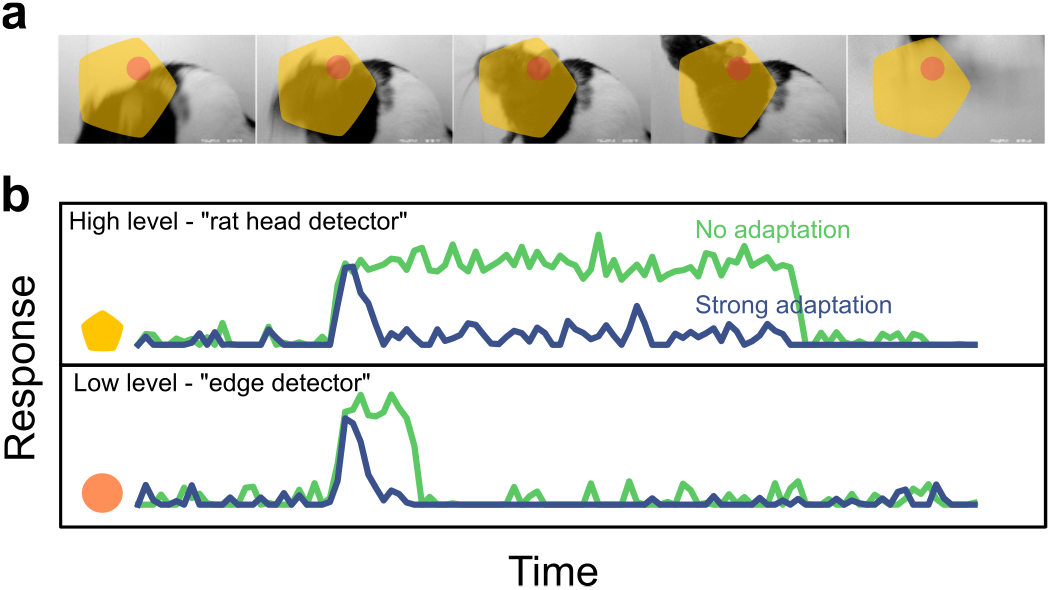
Effect of adaptation on the response timescale of high- vs low-level feature detectors (cartoon). **a** Dynamical visual stimulus (movie frames). Orange dot, yellow shape: idealized receptive field of a low-level feature detector neuron (“edge detector”, orange) and a high-level feature detector (“rat head detector”, yellow). **b** Single-trial response of the two example neurons, when adaptation is absent (green trace) and when adaptation is strong (blue trace). Note how adaptation shortens the timescale of the response.

An alternative scenario is one where mechanisms beyond history- and state-independent computations would significantly modify the temporal dynamics of neuronal responses, when tested with naturalistic dynamic inputs. For instance, adaptation has been shown to depress visual cortical responses to repeated presentation of perceptually similar visual stimuli in both monkeys and humans [21–23], resulting in a reduced discriminability of the adapted stimuli [24]. fMRI evidence in humans suggests that the impact of adaptation mechanisms may increase along the ventral stream [25–27] and that neural responses in higher areas of the hierarchy may be driven more strongly by transient than sustained stimuli [27; 28]. Similarly, in rats, adaptation has been shown to increase in magnitude along the cortical shape-processing hierarchy [29], attenuating the responses to predictable stimuli [30]. Finally, in primates, adaptation is at least partially preserved across object transformations (e.g., a smaller object can adapt the response to a larger object), as shown in both neurophysiological [31–33] and behavioral studies [34; 35]. Together, these effects could counteract any increase in response persistence resulting from increased invariance, as shapes that are temporally stable (e.g., a translating or expanding object) could strongly and continuously adapt neuronal responses until a new, surprising stimulus (i.e., a novel object) enters the neurons’ receptive fields. This is illustrated in Fig. 1b (blue curves), where the idealized “rat” detector unit, after an initial, transient response to its preferred object (the “rat head”), stops reporting the presence of the object within its RF, thus behaving similarly to a lower-level, edge-detector unit. More broadly, the predictive coding framework posits that, within certain cortical circuits, only error or surprise signals are encoded, their temporal dynamics naturally depending on a top-down signal carrying the input prediction [36–38]. This leads to the alternative hypothesis that the timescale and persistence of neural responses do not increase across the cortical hierarchy, as each cortical area encodes surprising features of the response of the previous area (see again the example cartoon in Fig. 1a, b). Some of these intuitions can be formalized by a computational model of adaptation [39], showing that response timescales to dynamic stimuli become progressively shorter as a function of the strength of adaptation (see Supplementary Text).

Yet other mechanisms exist that can alter the current state of the visual pathway in an activity-dependent way, but with qualitatively different effects than adaptation. For instance, temporally extended integration of synaptic input or recurrent excitation within local circuits can extend rather than shrink the temporal scale of visual processing [40; 41]. In presence of noise, their existence can be revealed by measuring the temporal span over which fluctuations of the firing rate are correlated (e.g., across repeated presentations of the same sensory input). This is typically called the intrinsic timescale of the recorded activity [42]. Interestingly, it was found that intrinsic timescales increase along various cortical hierarchies in primates and rodents [42; 43]. However, it is not known how the interaction between invariant encoding of object information and intrinsic processing unfolds along the ventral stream, and whether or not the net result is an ordered progression of temporal scales of neural processing. In addition, it is unknown how these processes interact with top-down modulation signals that reflect the attentive, motivational or locomotory state of a subject [44–49] and whose influence on the processing of dynamic visual stimuli is still largely unexplored.

To address the questions outlined in the previous paragraphs, it is necessary to perform neurophysiological recordings along an object-processing hierarchy during presentation of movie stimuli, ideally from the bottom to the top stage of the processing chain. The primate ventral stream would be the ideal target for such investigation, but the anatomy and size of the monkey brain makes it difficult to compare more than a pair of visual cortical areas in a single study (e.g., see [7; 8; 50]). By contrast, the small size of the rodent brain, the spatial contiguity of rodent visual cortical areas and the possibility to use a much larger number of animals than in monkey studies make it easier to probe and compare multiple visual regions (e.g., see [51–54]). Critically, during the last decade, a number of functional and anatomical studies in mice and rats have convincingly shown that visual information is processed in a hierarchical fashion across the many extrastriate visual areas that, in rodents, surround V1 [51; 52]. In particular, it has been shown that, along the progression of visual areas that run laterally to rat V1, many key tuning and coding properties evolve according to what is expected for a ventral-like, object-processing pathway [29; 30; 53–55], including the ability to support invariant visual object recognition – a finding that has been recently replicated also in mice [56].

Based on this wealth of evidence, we decided to compare the time scales of cortical processing of dynamic visual stimuli across the rat analogue of the ventral stream – i.e., V1 and the progression of extrastriate areas located laterally to V1: the lateromedial (LM), laterointermediate (LI) and laterolateral (LL) areas. To maximize the stability and duration of the recording sessions, as well as the number of repeated presentations of the movies used to probe these areas, neuronal recordings were performed under anesthesia [55; 57; 58]. We found that neural activity displays a hierarchy of temporal scales, as expected for a processing pathway where invariance is built through a cascade of feedforward computations. At the same time, we found that intrinsic temporal scales also become longer, and intrinsic correlations become more important in determining the overall temporal persistency of the neural representations, as one progresses from primary visual cortex (V1) towards higher-level areas. The generality of these findings was checked by analyzing two additional, existing datasets where visual cortex of awake rodents was probed with natural movies [59; 60]. Interestingly, the trends found in our experiments were replicated (being even sharper) in awake, head-fixed mice that were running on a wheel, but they were much attenuated during epochs in which the animals remained still and were fully absent in awake, head-fixed rats that were body restrained.

## Results

To investigate the temporal structure of neural representations of dynamic visual scenes, we used 32-channel silicon probes to record from cortical areas V1, LM, LI and LL in anesthetized rats. These areas are arranged in an anatomical progression that allow reaching each of them sequentially, with a single diagonal penetration, and stop at a desired location, so as to distribute the recording sites across 2 or 3 adjacent regions (see the example session in Fig. 2a left, where the probe targeted LI and LL).

**Fig. 2.**
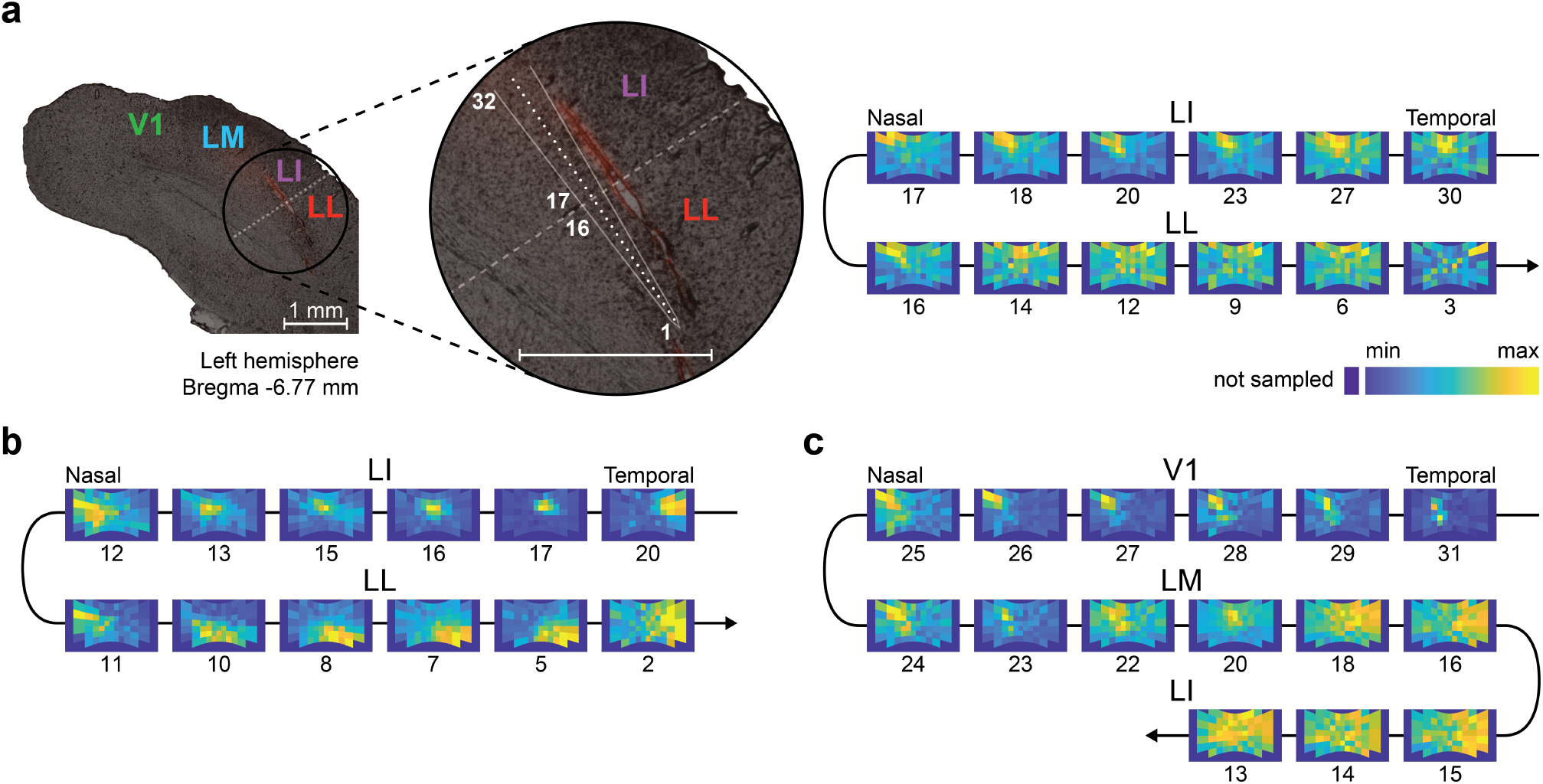
Functional identification of rat visual cortical areas. **a** Left: overlay of a fluorescence image on a bright-field image of a coronal slice of rat visual cortex, from which neuronal recordings were obtained using a single-shank silicon probe with 32 recording sites, spanning 1550 μm. Before being inserted obliquely into cortex (to target lateral extrastriate areas LI and LL), the probe was coated with a red fluorescent dye that allowed reconstructing its insertion track (red stain). The close-up view shows the geometry of the probe (white lines) alongside the insertion track and the relative positions of the recording sites (white dots), from the tip (site # 1) to the base (site # 32) of the probe. Right: firing intensity maps showing the RFs of the units recorded at selected recording sites along the probe (indicated by the numbers under the RF maps). The reversal in the progression of the retinotopy between sites 16 and 17 marks the boundary between areas LI and LL (shown by a dashed line on the Left panel). **b**, **c** Retinotopic progressions of the RFs recorded in two other example sessions. One session featured a single reversal between LL and LI within a span of the recording sites of 1550 μm (**b**). The other session featured two reversals, one between V1 and LM and another one between LM and LI, which were made possible by the larger depth spanned by the probe (3100 μm).

To infer the cortical area each unit was sampled from, we used a functional area identification procedure that has become the standard in electrophysiology and imaging studies of rodent visual cortex [51; 52]. That is, we tracked the progression of the RFs recorded along the probe, so as to map the reversals of the retinotopy that delineate the borders between adjacent areas [53–55]. To this aim, before the main stimulus presentation protocol with the movies, we run a RF mapping procedure, consisting in displaying drifting bars with four different orientations over a grid of visual field locations. The outcome of this procedure is illustrated, for an example recording session, in Fig. 2a (right), where the RFs recorded along the linear probe shown on the left are displayed as firing intensity maps over the visual field spanned by the monitor. Tracking this progression from the base (channel 32) to the tip (channel 1) of the probe revealed two distinctive retinotopic patterns: the RF initially shifted from the temporal to the nasal side of the visual field (channels 30-17), then headed back towards the temporal visual field (channels 16-3). This reversal of the retinotopy matched that reported in the literature for areas LI and LL [53–55; 61]. Another example of the reversal encountered while transitioning from LI to LL is shown, for a different session, in Fig. 2b. Fig. 2c shows instead an example session where the probe spanned three different areas (V1, LM and LI), resulting in two reversals of the retinotopy.

In each recording session, the RF mapping procedure was followed by the presentation of nine 20s-long movies, which included six natural dynamic scenes, as well as three synthetic movies – namely, a random sequence of white noise patterns and the Fourier phase-scrambled versions of two of the natural movies (see Methods and supplementary videos). Four of the natural movies (referred to as *manual* in what follows) were obtained by sweeping a hand-held camera at two different speeds (“slow” and “fast”) across an arena filled with various 3D-printed geometrical shapes that were painted either black or white (see example frames from one of the movies in Fig. 3a). The other two movies (referred to as *ratcam* in what follows) were obtained by placing a small camera on the head of a rat that was let free to explore the arena with the 3D-printed objects and another rat inside. Overall, a total of 294 well-isolated single units recorded in 18 rats were used in our analysis (see Methods for the selection criteria): 168 from V1, 20 from LM, 36 from LI, and 70 from LL.

**Fig. 3.**
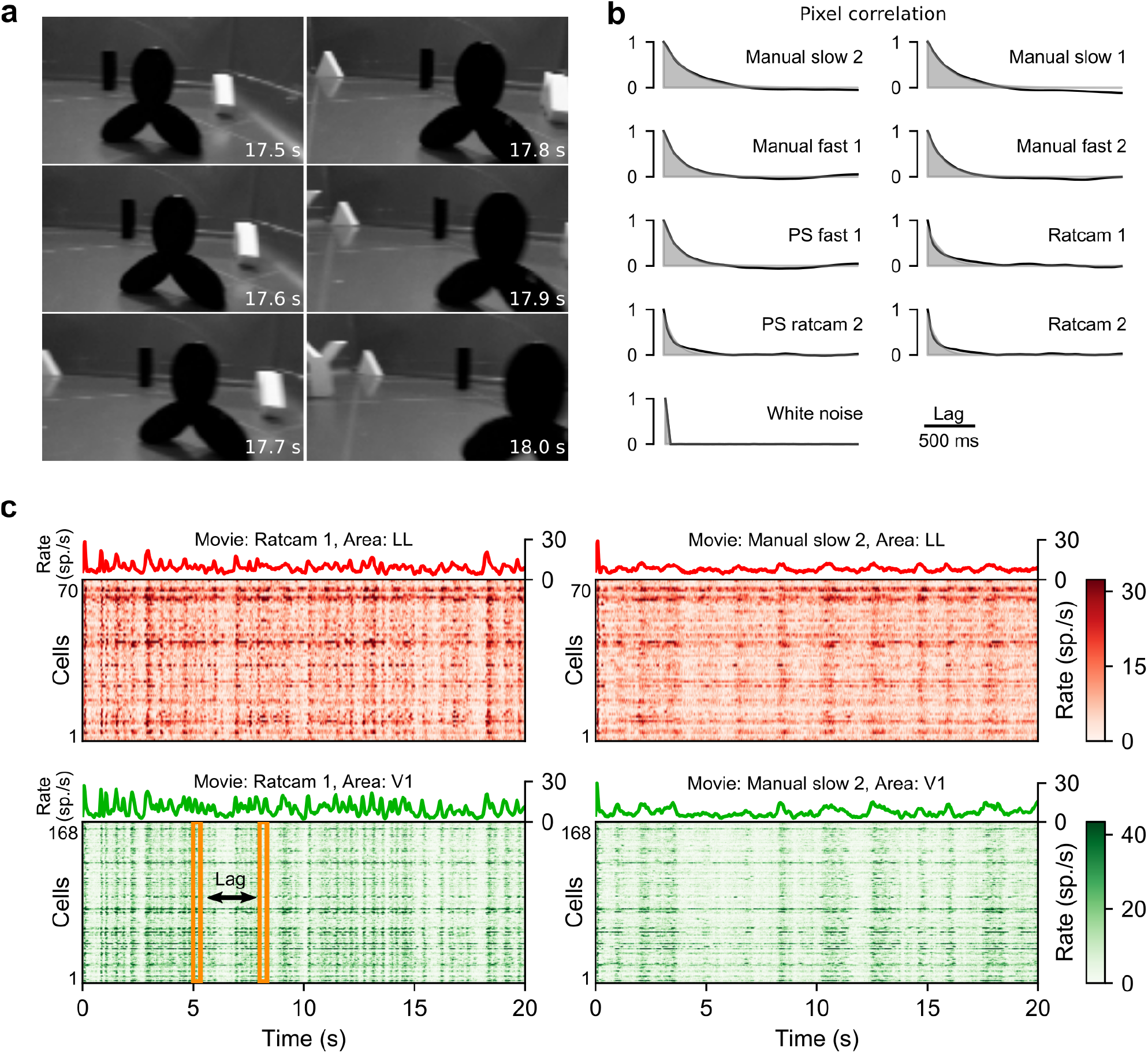
Visual stimuli and example neuronal recordings. **a** Six consecutive example frames from one of the movies used as visual stimuli in our study. The numbers on the bottom right of each panel give the time at which each frame appears. **b** Quantification of the characteristic timescale of the individual visual stimuli (movies). The *Manual* and *Ratcam* labels refer, respectively, to movies that were taken either by moving a hand-held camera across an arena filled with 3D-printed objects (some of them visible in **a**), or by installing a small camera over the head of a rat that was allowed to roam inside an arena containing the 3D-printed objects and another rat. The *PS* label refers to Fourier phase-scrambled versions of the corresponding movies (see Methods for details). In each panel, the black line shows the average correlation of pairs of image frames within a movie as a function of their temporal lag, while the shaded area shows the best exponential fit. **c** Examples of population responses to two of the movie stimuli. In the color matrices, each row color-codes the intensity of the trial-averaged response (computed in 33 ms time bins) of a neuron belonging to either the LL (red) or V1 (green) population. The color code is truncated to the 98^th^ percentile of the firing rate distribution within each area for ease of display. The trace on top of each matrix shows the corresponding population-averaged response. The yellow frames illustrate the procedure to compute the characteristic timescale over which the population activity unfolded, in response to a given movie (see Fig. 4a, b).

### Cortical representations unfold over timescales that are both stimulus- and area-dependent

As a first step in our analysis, we adapted an approach from [62] to characterize the temporal structure of the stimuli. Briefly, for each movie we computed the correlation coefficient between pixels belonging to image frames separated by a given lag. We then fitted the resulting dependence of the average correlation on the lag (Fig. 3b, solid line) with a decaying exponential (shaded areas). The time constant of this exponential defined the timescale of the movie (see Methods for details). According to this analysis, the dynamics of our stimuli spanned a range of timescales, from ~30 ms for the white noise movie to ~600 ms for the slowest of the natural movies.

The activity in each population was stimulus-modulated, in a consistent way across the recorded neurons, for both fast and slow movies. This is shown in Fig. 3c, where each line color-codes the response intensity of a LL (red) or V1 (green) unit to two example movies. As a result, the overall population-averaged activity (green and red traces) was also strongly stimulus modulated. To characterize how the temporal structure of the stimulus-locked responses depended on the visual input, we measured the population response timescale for each visual area and each movie by applying the same metrics defined above for the movie stimuli to the time-binned (bin width: 33ms) and trial-averaged population response vectors [62]. That is, with reference to Fig. 3c, we computed correlation coefficients between vertical slices (bins of spike counts) that were separated by a given lag (yellow frames), and then averaged the resulting coefficients to obtain a measure of signal correlation [63] as a function of lag (Fig. 4a, solid lines; in the following, we will call this type of signal correlations “response correlations”). This curve was fitted with an exponentially damped oscillation or a simple exponential based on a systematic model selection procedure (dashed lines; see Methods for details), and the decay constant of the exponential envelope was taken as the timescale of the response modulation. Next, we regressed the characteristic timescale of the response against the characteristic timescale of the stimulus, separately for each area, using a linear model with a common slope across all areas and an area-dependent intercept (Fig. 4b). The intercept is the baseline temporal timescale for stimulus-driven responses in each area. This baseline was clearly higher in the extrastriate areas (LM, LI, LL) than in V1, with smaller, statistically insignificant differences between LM, LI and LL (*p*=0.4 for both LM vs LI, and LI vs LL; two-tailed t-test, t=0.8,-0.8, df=31). Therefore, we repeated the regression analysis after lumping together these three areas (gray line). This revealed that the response timescales in all the areas depended strongly on the stimulus timescale (slope: 0.71±0.14, significantly different from 0 at *p*=1e-5, two-tailed t-test, t=5.2, df=33), and that the response timescale was significantly longer in extrastriate cortex than in V1 (intercept difference: 56±22ms; p=0.015, two-tailed t-test, t=2.6, df=33). Overall, these results show that the characteristic temporal scale of the population representation of dynamic stimuli depends on the temporal scale of the visual input, and that representations in extrastriate cortex unfold over longer temporal scales than in V1.

**Fig. 4.**
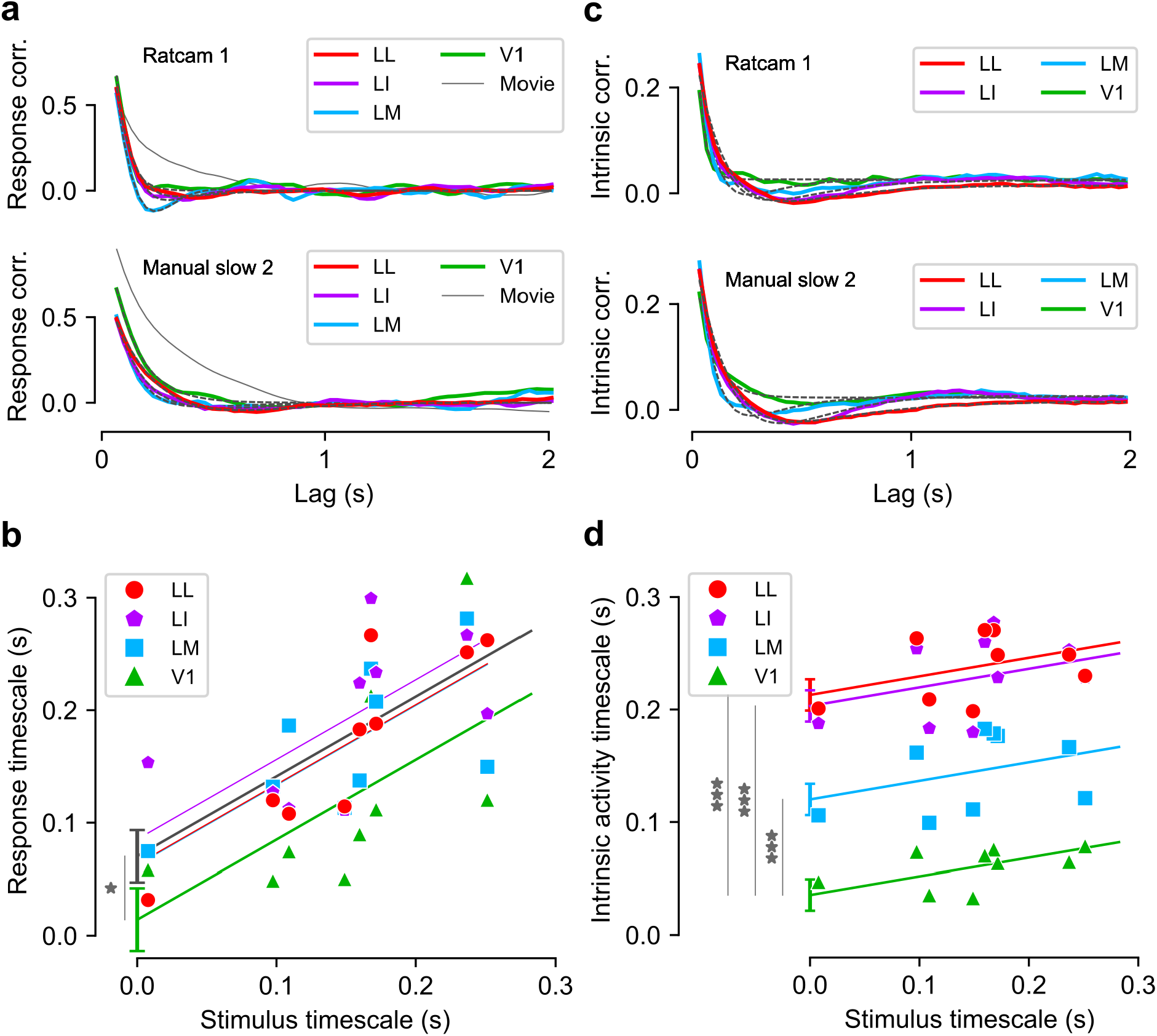
Characteristic timescale of the stimulus-driven population responses and of the intrinsic neuronal activity. **a** Correlation between pairs of population response vectors (as those highlighted by the yellow frames in Fig. 3c) as a function of their temporal lag for two example movies (solid, colored lines). These curves were fitted with either an exponential decay or a damped oscillation function (fits are shown as black dashed lines). The time constants of these fits were taken as the characteristic timescales of the population responses. The solid gray lines show the correlation functions of the corresponding movies, reported from Fig. 3b for comparison. **b** Timescales of the population responses as a function of the timescales of the corresponding movie stimuli (colored markers). Each colored line is the linear fit prediction of the relationship between such timescales for a given area (same color code as in the key; note that the lines for LM and LL are partially overlapping). The gray line is the linear fit prediction obtained by pooling together the data of the three extrastriate areas (i.e., LM, LI and LL). The error bars show the standard errors of the intercept of the linear fits for V1 and the pooled extrastriate areas. **c, d** Same as **a** and **b**, but for the intrinsic timescales of neuronal activity (see main text for details).

### The timescale of intrinsic processing increases along the ventral-like pathway

As illustrated in the previous section, our results suggest that the timescale of stimulus-locked, trial-averaged neural representations increases from V1 to extrastriate cortex. The temporal scale of intrinsic neural processing has also been suggested to increase across cortical hierarchies at the single cell [42] and at the population level [43]. We tested whether this applied to the progression of rat lateral visual areas by using the method described by [42], which is mathematically similar to the procedure used above to compute the timescale of the population response vectors, but considers the responses of a single cell across multiple trials, rather than the average responses of multiple neurons (see Methods and Supplementary Fig. 1). This allowed us to capture the largely stimulus-independent, within-trial temporal correlations in the spiking activity of a neuron, which were then averaged over all the units of a population. The time-dependence of the resulting correlation function (solid lines in Fig. 4c) was fitted with an exponential or an exponentially damped oscillation, based on systematic model selection performed independently for each condition (black dashed lines; see Methods for details). The characteristic temporal scale was determined as the decay time constant of the exponential envelope of the fit. Finally, we linearly regressed the intrinsic timescale of neural activity with the timescale of the movie stimulus and the cortical area (Fig. 4d), as we did above for the stimulus driven response (Fig. 4b). The intercept values of the linear fit revealed a clear hierarchical organization of the timescale of intrinsic activity (V1: 36±14ms; LM: 120±14ms; LI: 203±14ms; LL: 213±14ms), with all three extrastriate areas being significantly different from V1 (*p*=5e-7, 1e-13, 2e-14 respectively for LM, LI, LL, two-tailed t-test, t=6.3, 12.6, 13.3, df=31). We also found a mild dependence on the timescale of the movie (slope: 0.17±0.07, *p*=0.02, two-tailed t-test, t=2.4, df=31), much weaker than that observed for the stimulus-driven response (compare Figs. 3b and d). Interestingly, the range of intrinsic timescales we recorded in our experiment quantitatively matched that reported for sensory cortex in behaving monkeys by [42] and in behaving mice by [43]. Overall, these results show that the temporal scale of the intrinsic activity increases along rat lateral extrastriate visual areas.

### The temporal stability of neural representations increases along the ventral-like hierarchy

Temporal stability of a neuronal representation supports stable perception of the content of a visual input that unfolds smoothly over time. For instance, if an object (e.g., a triangle) sweeps across the visual field, a temporally stable representation should support discrimination of this object from other objects (e.g., squares) despite ongoing continuous changes in its appearance. Correlation between neural activity at different times does not by itself assess such stability of discrimination because population firing vectors can have a fixed amount of correlation and yet be discriminable to different degrees, depending on how they are organized with respect to a discrimination boundary. Moreover, a correlation measure assumes that deviations in the neural code are most important along directions that are orthogonal to the current population vector, as correlation coefficients behave like a dot product or a cosine similarity measure. This is not necessarily the right notion of similarity for discrimination problems. Thus, we sought a direct test of the temporal stability of discrimination based on neural population responses.

To this end, we imagined a task where an observer learns to discriminate between the visual input appearing at a time *t*_*A*_ and that presented at a much later time *t*_*B*_. If this task is performed on the basis of the activity in a given cortical area, we should be able to train a classifier to reliably discriminate population response vectors occurring at these two times in the movie. Now consider responses at a pair of time points shifted by a small lag *Δt*, namely *t*_*A*_ + *Δt* and *t*_*B*_ + *Δt*. If the relevant part of the representation of the visual input is temporally stable over this lag, the trained classifier should perform as well on responses at the lagged time points as it did at the original time points (see the schematic in Fig. 5a). Averaging over all *t*_*A*_ and *t*_*B*_ for each lag *Δt* will assay how well the population responses support temporally stable discrimination of visual inputs.

**Fig. 5.**
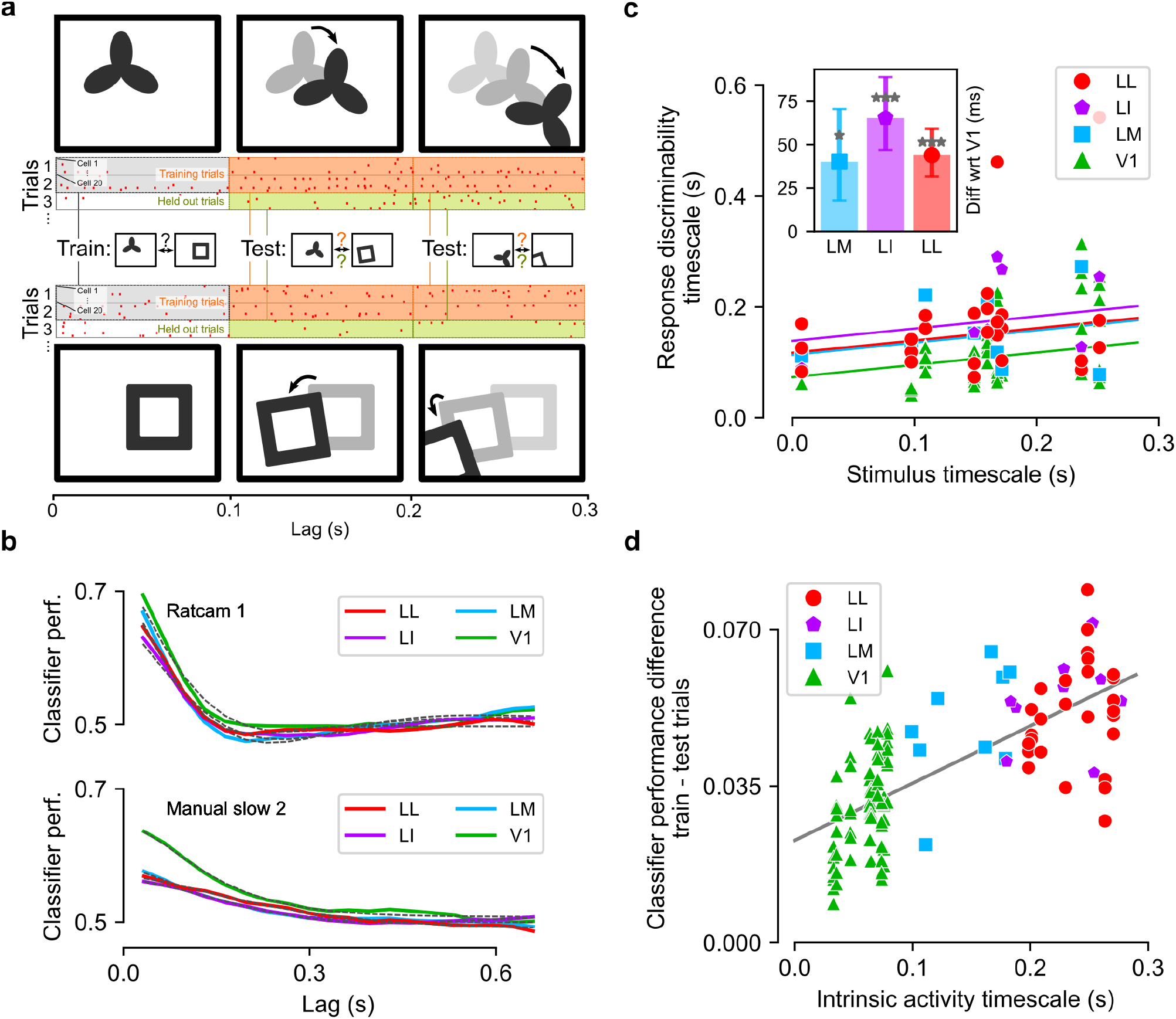
Temporal stability of neuronal representations, as assessed through decoding analysis. **a** Schematic of the classifier analysis, aimed at assessing the temporal stability of the population representation. A linear classifier is trained to distinguish the population activity recorded at two different time points of the movie (spike counts computed from the gray time bins) and tested at other time points that follow (or precede, not shown) the training points by a certain lag. The classifier is tested on a held-out set of trials (green boxes), as well as on the same experimental trials that were used for training (orange boxes). The cartoons of the movie frames shown on top or below each time bin are for illustration purpose only: they highlight the fact that, in general, two non-overlapping segments of a movie (from which the population activity is decoded) will contain different, transforming (e.g., translating, rotating, etc.) objects. For clarity, in this illustration, the size of the spike count windows (i.e., gray, orange and green boxes) was set to 100 ms, while 33 ms wide bins were used in the actual analysis. The spike patterns shown in the cartoon correspond to those actually fired by a randomly selected pseudopopulation of 20 units in V1 at two time points during the presentation of one of the movies (only three trials shown out of 30). **b** Classifier performance on held-out trials for two example movie stimuli and one example pseudopopulation per area (colored curves). The performance is plotted as a function of the lag between training bin (gray boxes in **a**) and test bin (green boxes in a), and is fitted with either an exponential decay or a damped oscillation function (fits are shown as black dashed lines). **c** Timescale of response discriminability, measured as the time constant of the exponential decay of the classifier performance on the held-out trial set, as a function of the timescale of the movies. Each dot corresponds to a distinct pseudopopulation of 20 units. The solid lines are linear regressions with common slope and different intercept across the four areas. The inset shows the difference of the intercept for areas LM, LI and LL vs. area V1 (* *p* < 0.05, *** *p* < 0.001, one-tailed bootstrap test). **d** Amount of classifier performance due to intrinsic correlations (Δ_*p*_, Eq. 1) as a function of the timescale of such correlations (i.e., the intrinsic timescale of neuronal activity shown in Fig. 4d).

Following this strategy, we implemented a linear classifier on pseudopopulations of 20 randomly selected units (see Methods). Results for larger populations (50 units) from areas where these were available (V1 and LL) are reported in Supplementary Fig. 2. The classifier was trained to distinguish population activities, represented as 20-dimensional spike count vectors (bin width 33ms), evoked by a movie at two different time points. Critically, only a subset of trials was used for training (gray boxes in Fig. 5a). The temporal stability of the population code was then assessed by testing the performance of the decoder at time points shifted by a lag *Δt* (Fig. 5a). In order to isolate the contribution of the direct drive from the visual stimulus to the overall temporal stability, we tested the classifier only on held-out experimental trials that were not used in training (green boxes). This ensured that intrinsic temporal correlations (Fig. 4d) between lagged frames would not affect the similarity between activity vectors used in training and testing. This analysis was repeated for each population and each movie, and for all possible choices of pairs of training time points (details in Methods). The results were averaged over these pairs to measure the mean discrimination performance as a function of the lag (Fig. 5b; colored lines), and then fitted with the same functional form used above for the temporal correlations of the neural responses (black dashed lines). A linear regression revealed that the decoder’s performance depended significantly on the timescale of the movie stimulus (Fig. 5c; common slope: 0.22, 68%CI [0.15-0.32], nonzero at *p*=8e-4, one-tailed bootstrap test). In addition, the intercept of the fit was significantly larger in the three extrastriate areas than in V1 (V1 intercept: 74[55-82]ms; difference with respect to V1 for the other areas: LM 40[17-70]ms, nonnegative *p*=0.03; LI 65[48-89]ms, *p*<1e-4, LL 44[32-59]ms, *p*=2e-4, one-tailed bootstrap test). Such a difference indicates that neural representations in higher areas of the ventral hierarchy support percepts of dynamic stimuli that are stable over longer timescales as compared to V1 representations.

### Intrinsic correlations extend the within-trial temporal stability of neural representations

As mentioned in the Introduction, intrinsic correlations among fluctuations of neuronal firing at different times can be interpreted as the tendency of a neuron to sustain a given firing rate level over time, regardless of the incoming sensory input. As such, intrinsic correlations have been argued to be the hallmark of longer stimulus integration time, or to act as “stabilizers” of neuronal representations to better support reading of the latter by downstream decision areas [42; 43; 64]. In the vision domain, object perception unfolds continuously on the basis of the ongoing activity in neural populations of visual cortical areas. Intrinsic temporal correlations in this activity (Fig. 4d) could stabilize the neural code against changes in the visual input, maintaining a memory trace of the objects moving across the visual field, thus stabilizing their perception.

To test for this possibility, we repeated the classification analysis described in the previous section. This time, however, we measured the performance of the trained classifier at lagged time points in the same trials that were used for training (orange boxes in Fig. 5a; the resulting performance curves are shown in Supplementary Fig. 3a). In this case, fluctuations in test response vectors at times *t*_*A*_ + *Δt* and *t*_*B*_ + *Δt* will be correlated with fluctuations in the training vectors at *t*_*A*_ and *t*_*B*_. In fact, a neuron responding to a given movie frame, in a given trial, with a number of spikes larger (or smaller) than its mean response across trials, will tend to over-fire (or under-fire) also in the following frames within the same trial. To quantify the contribution of these intrinsic correlations to the stability of discrimination we focused on the performance of the classifier at a lag of 33ms (one video frame) from the training bin, and we computed the gain of performance due to intrinsic correlation as follows:

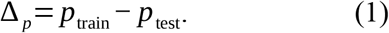

Here *p*_*train*_ and *p*_*test*_ are the performances of the classifier at lag 33ms, when evaluated on the training trials (orange boxes in Fig. 5a) or on the held-out trials (green boxes), respectively.

As expected, given the progressively longer intrinsic correlations observed along the cortical hierarchy (see Fig. 4d), *Δ*_*p*_ substantially increased from V1 to LL (Fig. 5d. Slope of the fit: 0.13±0.01, nonzero *p*=*2e-18*, two-tailed t-test, t=10.5, df=115; intercept: 0.023±0.002, p=6e-24, two-tailed t-test, t=12.8, df=115), being, on average, almost twice as large in the latter than in the former (i.e., 0.051 ± 0.002 in LL vs. 0.03 ± 0.001 in V1). This revealed that intrinsic correlations play an increasingly important role in determining the overall temporal stability of the neural code along the cortical progression. Overall, these findings show that the relative importance of intrinsic dynamics for population codes in visual cortex increases along a ventral-like, object-processing hierarchy.

### Temporal stability as a signature of invariance of neural representations

The decoding analysis shown in Fig. 5 indicates that representations in rat lateral extrastriate areas remain stable over longer timescales than in V1. However, in terms of magnitude of the decoding performance, V1 was found to surpass all other areas, at least at time lags that are close to the training bin (see Fig. 5b). In other words, the increase of invariance that is expected to take place along an object-processing hierarchy was observable only in relative terms (i.e., decoding performances changing less over time in LM, LI and LL), rather than in absolute terms (i.e., decoding performance being larger in LM, LI and LL). This result can be understood on the basis of our previous study of rat lateral extrastriate cortex using static objects [53], and by considering the movie segments that were used to carry out the decoding analysis in Fig. 5.

[53] showed that LI and LL representations do afford larger decoding performances of object identity than V1 and LM representations under a variety of transformations in object appearance (i.e., position, size, rotation, etc.). However, the larger invariance of higher-order vs. lower-order areas only emerged when the pairs of objects to discriminate had similar levels of luminosity across the tested transformations. Otherwise, the objects were actually better discriminable based on the V1 representation than the one in LL. This finding can be explained based on another result of our previous study – the information conveyed by neuronal responses about low-level visual properties, such as luminosity and contrast, decreases substantially from V1 to LL. Such pruning of low-level information is fully consistent with ventral-like processing, but has the unexpected effect of making it easier for V1 neurons to discriminate visual objects in case their average luminance (across transformations) is not matched. As explained in Tafazoli et al (2017), the superior invariance observed for V1 under such scenario is only apparent, “because luminance would not at all be diagnostic of object identity, if [V1] neurons were probed with a variety of object appearances as large as the one experienced during natural vision”.

Given the nature of the stimulus set used in [53] (bright, isolated shapes against a black background), it was possible there to restrict the decoding analysis to object pairs with similar luminance, thus revealing the larger invariance of LL. This was not possible with our current, more naturalistic stimulus set. As a result, differences in luminance and other low-level properties likely played a major role in allowing the V1 representation to afford the largest decoding performance. In fact, given the continuous nature of our dynamic stimuli, luminance and contrast differences between the movie frames used to train a classifier would be preserved in the preceding and following frames. They would only vanish at long time lags from the training bins. Hence, the larger decoding performance achieved by V1 at short time lags in Fig. 5b.

To verify the correctness of this interpretation, we repeated our decoding analysis after selecting 21 movie segments with identical duration (967 ms, or 29 movies frames) that contained a single object (either black or white) moving roughly horizontally, either from left to right or from right to left (see examples in Fig. 6a). This allowed defining four different types of classification tasks, based on whether the color and/or motion direction of the objects to discriminate were (or not) the same (Fig. 6a, b). Our expectation was that, at least for tasks where the difference in object luminance was small, extrastriate areas would reach the same absolute performance level as V1 and possibly surpass it at sufficiently long lags from the training bin. In particular, we expected this to be the case when the objects in the movie segments moved along opposite directions. In this case, the large sensitivity to stimulus position of V1 neurons [53; 65] would prevent them to keep track of object identity at long time lags, when the two objects have fully swapped their position as compared to the training bin.

**Fig. 6.**
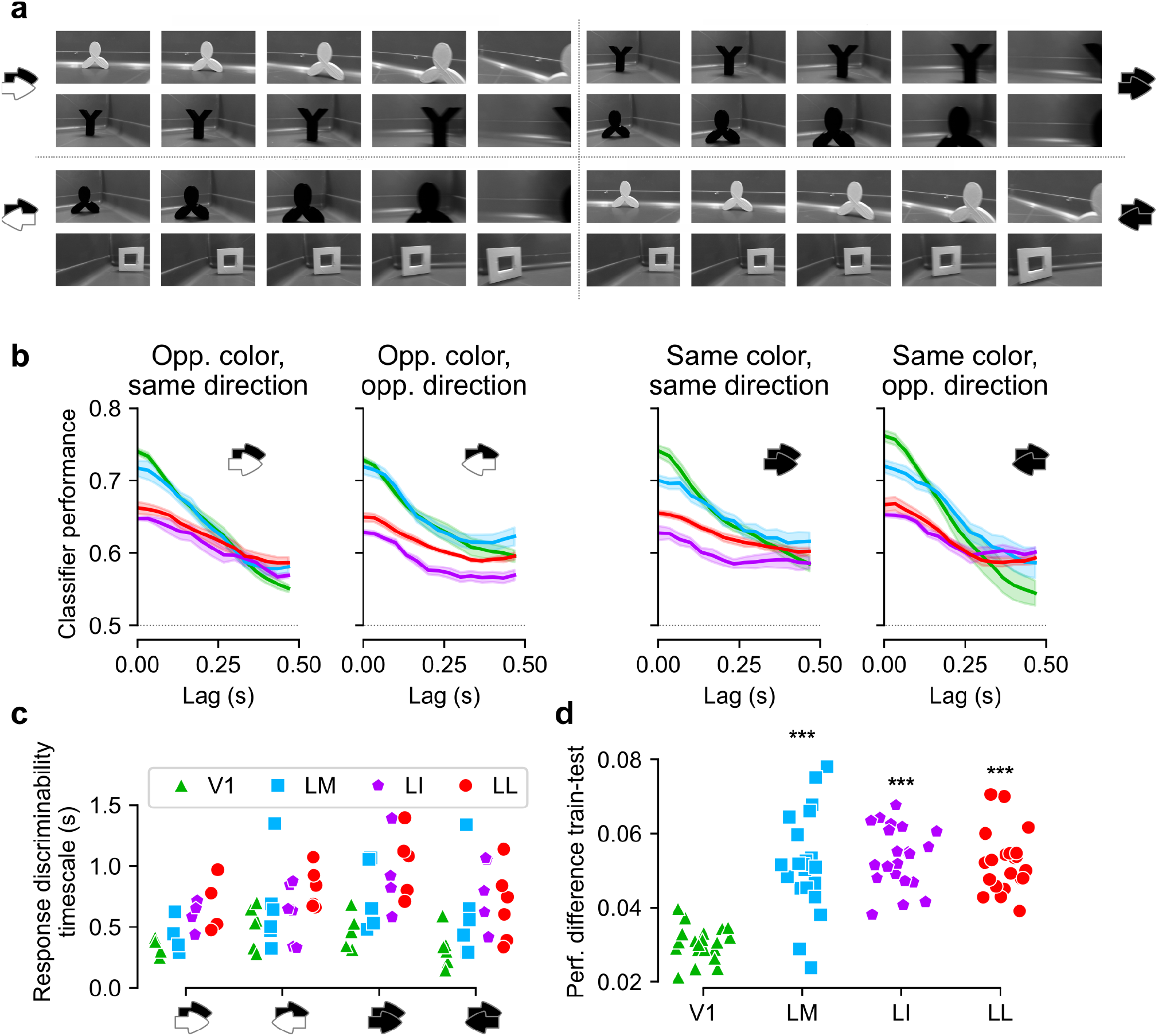
Classifier analysis restricted to hand-matched pairs of movie segments. **a** Frames of four example pairs of movie segments (each pair is representative of one of the four classification conditions that were tested in our analysis). Each movie segment was exactly 29 frames long, and the frames shown here are frames number 1, 8, 15, 22, and 29 (left to right). As indicated by the black/white arrows in each quadrant, each pair of movie segments was distinguished from the others, based on whether the objects in the segments had the same color and whether they moved along the same direction. **b** Classifier performance vs. lag from the training bin for each of the four conditions. Solid line: average over all pairs tested in the task. Shaded area: s.e.m. **c** Timescales extracted from the decoding performances of all pairs of movies segments tested in any given classification task and for each area (two outlier data points are not shown: 3.65s for same color/same direction, and 2.73s for same color/opposite direction). **d** Amount of classifier performance due to intrinsic correlations (difference between performance on training trials and on test trials; as defined in Fig. 5d) for each movie pair and each area. Significance markers: *p* < 0.001, paired t-test on each area vs V1.

This new decoding analysis also allowed measuring the stability of visual representations in a scenario where object identity did not to change within a movie segment. By contrast, in our previous analysis (Fig. 5), the segments fed to the classifiers were from all possible locations along the whole movie. Thus, every segment could contain multiple objects, and these could appear and disappear (being replaced by other objects) while the movie unfolded over time (e.g., see Fig. 3a). This analysis is ideal to obtain an overall, unbiased assessment of the stability of cortical representations of a given movie. However, it does not allow measuring the temporal persistence of the representations of individual objects undergoing identity-preserving transformations. It is the latter kind of stability that is expected to be equivalent to the invariance measured with static objects and, as such, is expected to grow more along an object-processing pathway. Therefore, the time constants of the decoding curves shown in Fig. 5 are likely an underestimation of the actual persistence (or invariance) of object representations in the four visual areas, and the larger stability of extrastriate areas over V1 is likely also underestimated.

The results of our new decoding analysis (Fig. 6b) largely confirmed these intuitions. As expected, in terms of magnitude of the decoding performance, V1 surpassed all other areas at short time lags. However, the performance afforded by V1 neurons decayed very abruptly over time, being matched and even surpassed by that yielded by the other areas at longer time lags. Most notably, when the objects had the same color but opposite direction (rightmost panel), the larger position-invariance of the extrastriate areas emerged very clearly, with the V1 performance dropping substantially below that of the other areas at time lags > 300 ms. This confirmed the prediction that in a task very demanding in terms of invariance (i.e., objects swapping position across the visual field) and less prone to be solved based on luminance differences (i.e., object with the same color), lateral extrastriate areas would have emerged as the most invariant [53].

As expected, restricting the decoding analysis to movie segments with single objects also resulted in a much slower dynamics (compare the performance curves of Fig. 6b to those of Fig. 5b). This increased temporal stability of object representations was particularly strong for high-order extrastriate areas (LI and LL), and, to a lesser extent, for LM. In V1 instead, the representation unfolded over a much faster timescale. Fitting exponentially damped oscillations to the performance curves yielded time constants (Fig. 6c) that, in LI and LL, were up to five times larger than those obtained in our previous, unconstrained analysis (Fig. 5c). More importantly, the time constants in LI and LL were about twice as large as in V1, thus revealing a substantially slower unfolding of higher-order representations. A formal statistical test based on categorical linear regression (see Methods) confirmed that the time constants in LI and LL were higher than in V1 (timescale difference with respect to V1: LM, 210 ± 140 ms; LI, 550 ± 140 ms; LL, 390 ± 140 ms; p-values, two-tailed t-test: LM 0.13, LI 0.0001, LL 0.006).

In the analysis of Fig. 6B,c, as previously done in Fig. 5b, c, the generalization ability of the classifiers was tested on held-out trials that were not used for training. To gauge now the impact of intrinsic temporal correlations in stabilizing object representations, we measured generalization at lagged time points in the same trials that were used for training (same kind of analysis previously shown in Fig. 5d). As before, the increment of performance *Δ*_*p*_ (see eq. 1) afforded by intrinsic correlations was substantially and significantly higher in lateral extrastriate areas than in V1 (Fig. 6d; mean *Δ*_*p*_: V1, 0.030 ± 0.001; LM, 0.052 ± 0.003; LI, 0.054 ± 0.002; LL, 0.052 ± 0.002; paired t-test p-values for the difference with respect to V1: LM p=1e-7, LI 7e-10, LL 3e-9). This confirmed the increasingly important role played by intrinsic correlations in determining the overall temporal stability of object representations along the cortical progression.

### Temporal stability of visual cortical representations in awake mice

The results presented in the previous sections are based on recordings performed in anesthetized rats. To validate them in awake animals, we repeated our analysis on a large dataset of neuronal responses recorded from many visual areas of awake mice using Neuropixels probes [60] – a dataset that was recently made freely available by the Allen Institute for Brain Science as part of the Allen Brain Observatory. The fact that the recordings were performed on mice added the advantage of testing the generality of our conclusions on a different rodent species. Another advantage is that the ranking of mouse visual areas in terms of hierarchical processing has been carefully established at the anatomical level [67; 68] and recently quantified by the definition of an anatomical hierarchy score [66]. This score can be related to a number of functional measures of hierarchical processing, thus checking the consistency between anatomy and physiology [60].

Following this approach, we tried to establish whether, in the Allen dataset, the temporal stability of neuronal responses to dynamic stimuli increased as a function of the hierarchy score of mouse visual areas. To this aim, we focused on the two natural movie clips (one lasting 30 s and presented 20 times, the other lasting 120 s and presented 10 times) that were part of the battery of visual stimuli tested in [60]. The shortest clip was discarded, given that its pixel correlations displayed a strongly irregular pattern (see Methods and Supplementary Fig. 4a, left), likely because the clip is composed by a sequence of highly static scenes. We thus carried out our analyses on the longest clip, whose pixel correlation function has an exponential decay (Supplementary Fig. 4b, green curves) that is fully comparable with that of our stimuli (see Fig. 3b). We extracted the neuronal responses to the repeated presentations (trials) of this movie that were recorded in the first 17 sessions of the Allen dataset from the following visual areas: the lateral geniculate nucleus (LGN) in the thalamus (571 cells), V1 (876 cells), and five higher-order, extrastriate visual cortical areas: LM, AL, RL, PM, and AM (506, 511, 796, 565 and 687 cells respectively). For each area, we then computed the correlation function of the population responses vectors (same analysis illustrated in Fig. 3c), as well as the population-averaged intrinsic correlation function, capturing within-trial fluctuations of the firing rate (same analysis illustrated in Supplementary. Fig. 1).

The resulting correlation functions are shown in Fig. 7a and b (solid lines), along with the best exponential fits to the data (dashed lines; the same fitting procedure used for the analysis of Fig. 4a and c was used; see Methods for more details). From these fits, we obtained the time constants of the exponential decays across the 7 areas and we regressed them against the corresponding hierarchy scores. For both the stimulus-driven (Fig. 7c) and the intrinsic (Fig. 7d) time constants we found a clear, significant linear relationship (respectively, p=0.001, t=5.8, and p=0.01, t=3.3; 1-tailed t-test, df=5). That is, we observed an increase of temporal stability across the visual anatomical hierarchy of awake mice that was very similar to that previously found across the object-processing pathway of anesthetized rats.

**Fig. 7.**
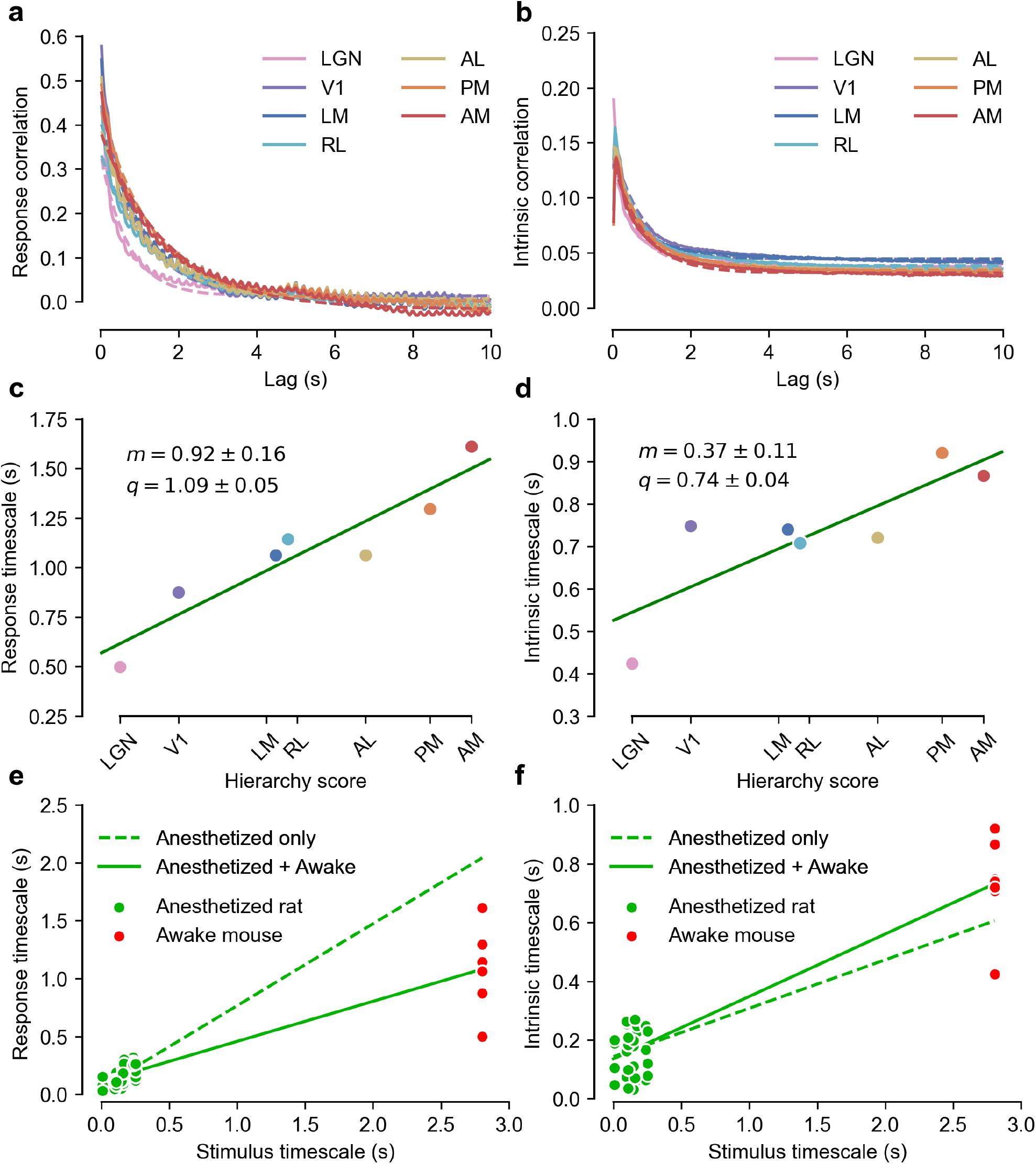
Timescale of stimulus-driven responses and intrinsic neuronal activity in awake mice (Allen dataset). **a** Correlations in the responses to a 120 s-long movie clip that were recorded across seven visual areas of the mouse visual system by [60] and are part of the Allen dataset. The same color code used by [60] denotes the different areas (see key). Dashed lines with matching colors indicate the corresponding exponential fits. **b** Same as in **a** but for the intrinsic correlations. **c** The time constants of the exponential fits to the correlation functions shown in **a** are plotted against the anatomical hierarchy score of the corresponding visual areas, as measured by [66]. m: slope of the best linear fit. q: intercept of the best linear fit. **d** Same as in **c** but for the intrinsic correlations shown in **b**. **e** All the response timescales (without area distinction) obtained in anesthetized rats (green dots; same data as in Fig. 4b) and in awake mice (red dots; same data as in **c**) are plotted against the corresponding stimulus timescales. The dashed green line shows the linear regression performed just on the rat anesthetized data, while the solid line shows the regression performed on the combined data. **f** Same as in **e** but for the intrinsic timescales previously shown in Fig. 4c for the anesthetized rat recordings (green dots) and in **d** for the awake mouse recordings (red dots).

On the whole, the time constants measured for the Allen dataset were substantially larger than those measured in our experiments. This is not surprising, given that the Allen movie clip unfolded over time with a much slower dynamics than the movies used in our recordings (compare Supplementary Fig. 4 to Fig. 3b) and, based on the trends reported in Fig. 4b and d, we expected the dynamics of the response to track that of the stimulus. As shown in Fig. 7e and f, the latter conclusion was confirmed when the time constants obtained in our recordings (green dots; same data of Fig. 4b and d) and those extracted from the Allen dataset (red dots; same data of Fig. 6c and d) were plotted against the time constants of the corresponding movies. The dependence between response and stimulus dynamics that could be inferred from our recordings extrapolated very well to the Allen dataset, with the red dots being very close to the values expected based on a linear regression (dashed green line) performed on the green dots only. This is striking, when considering how narrow the span of temporal scales of our movies was (0-0.25 s), with respect to the time constant of the Allen movie clip (2.8 s).

### Dependence of temporal stability on the behavioral state of the animal

The results of the previous section indicate that the conclusions of our study are highly robust with respect to the rodent species under exam (i.e., rat vs. mouse) and to the state of the animal (i.e., whether anesthetized or awake). However, the awake state can be further broken down into a spectrum of finer brain states, depending on the level of alertness, motivation, appetite or activity of the animal [44; 46–49]. In particular, recent studies have shown that locomotion increases response magnitude and spatial integration in mouse visual cortex [44; 46]. More importantly, during active wakefulness, mouse visual cortical representations of natural movies have been found to be more similar to those measured under anesthesia than during quiet wakefulness [47].

Motivated by these previous findings, we further explored the Allen dataset, given that mice, during recordings, were head-fixed but they were free to either rest or run on a spinning wheel, whose angular velocity was part of the acquired data. Based on this information, we identified epochs of the neuronal responses to the movies during which the mouse was either resting or running (see Methods for details). We then computed the response and intrinsic timescales separately for the resting and running states. In general, the observed trends slightly changed depending on the number of trials *N*_*t*_ (i.e., stimulus repetitions) included in the analysis and on the minimal duration *L* of the resting (or running) epochs that were shared among the chosen trials. Given that these two parameters were inversely related (see Methods and Supplementary Fig. 5), we considered all combinations of four choices of *N*_*t*_ (i.e., 3, 4, 5 and 6 trials) and five choices of *L* (i.e., 2.5, 3.3, 4.1, 5.0 and 6.6 s).

Overall, we found that the increase of temporal stability observed across the mouse visual hierarchy was weaker in the resting than in the running state. This is illustrated for an example choice of the parameters *N*_*t*_ and *L*, in Fig. 8 (leftmost plots), which shows how the slope of the linear fit through the time constants was smaller in the case of the resting state, especially for the intrinsic dynamics (Fig. 8b). As shown in Fig. 8a (right), in the case of the response timescales, the reduction of the slope of the fit from the resting to the running state was observed systematically across most of the 4 × 5 combinations of the parameters *N*_*t*_ and *L*, overall yielding a significant difference (two-tailed, paired t-test; p=0.002, t=3.6, df=19). The effect was even stronger for the intrinsic timescales (Fig. 8b, right), where the slope of the fit decreased by more than half in the resting as compared to the running state across all tested combinations of the parameters *N*_*t*_ and *L*, thus yielding a highly significant difference (two-tailed, paired t-test; p=2e-8, t=9.2, df=19).

**Fig. 8.**
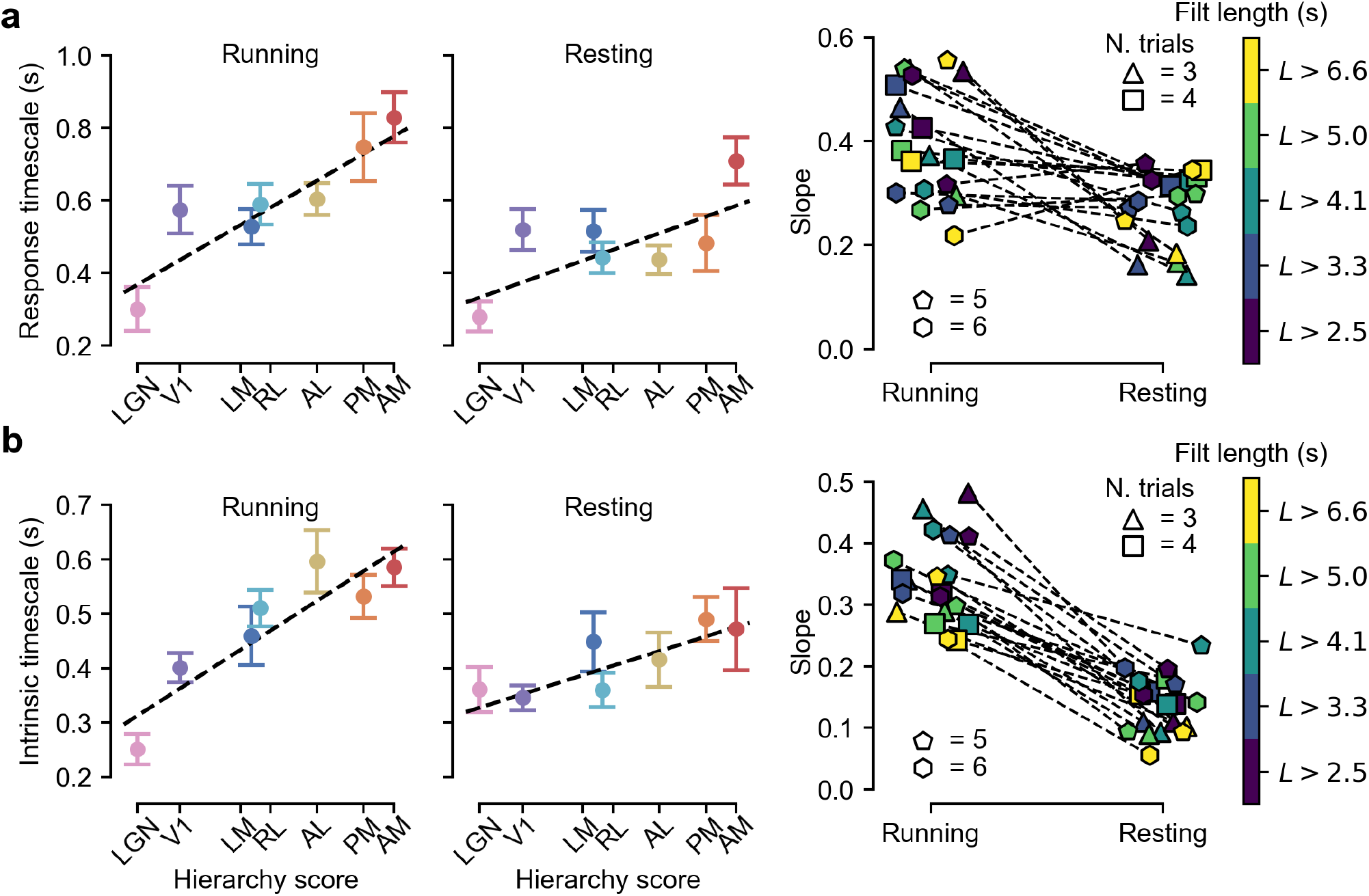
State dependence of response and intrinsic timescales. **a** Left, center: examples of the hierarchical trends of response timescales as a function of the anatomical hierarchy score for the particular choice of the parameters *N*_*t*_ = 5 and *L* = 4.125s (see main text) in the running (left) and resting (center) state. Data points are averages over 11 epochs of running and resting states. Error bars are s.e.m. The dashed black lines are error-weighted linear regressions. Right: scatter plot reporting the angular coefficients of the linear regressions for all the 4 × 5 combinations of the parameters *N*_*t*_ and *L*, in the case of the running and resting states (the dashed lines connect matching choices of *N*_*t*_ and *L*). The shape of the markers codes for *N*_*t*_, the color for *L* (see key). **b** Left, center: same as in **a** (left, center) but for the intrinsic timescales and for the choice of the parameters *N*_*t*_ = 6 and *L* = 2.475s. Data points are averages over 11 epochs of running and resting states. Right: same as in **a** (right) but for the intrinsic timescales.

Overall, this analysis shows that, during quiet wakefulness, the increase of temporal stability across the visual processing hierarchy is substantially reduced as compared to the state of active wakefulness, especially at the level of intrinsic correlations. To further investigate this phenomenon, we analyzed a third dataset of neuronal responses to natural movies that have been collected in head-fixed and body-restrained awake rats in the context of a previous study of the rat object-processing pathway [59]. In this study, neuronal responses were obtained from two of the regions also sampled in our recordings, i.e., V1 (44 cells) and LI (40 cells), plus a third visual area, named TO (38 cells), that follows LL in the anatomical progression of lateral extrastriate areas and that a previous study indicates to be part of the rat ventral-like pathway [54].

In this head-fixed rat dataset, the autocorrelations of the stimulus-driven responses displayed exponential decays with time constants that were not significantly larger in high-order areas (i.e., LI and TO) than in V1 (Supplementary Fig. 6a, b). In addition, the dependence between the temporal scale of the response and that of the movie was very weak and not significant (regression slope: 0.07 ± 0.07, *p* = 0.3, t = 1.04; intercept difference extrastriate-striate: −0.015 ± 0.020, *p* = 0.46, *t* = - 0.74; df = 57; see Methods for details on the regression). As for the intrinsic correlations, they were weaker than those found in our recordings and in the Allen data and did not display a monotonic decrease as a function of time (compare Supplementary Fig. 6c to Figs. 4c and 7b). We nevertheless managed to fit exponential decays to the tails of these autocorrelation functions. The resulting time constants did not increase from V1 to higher-order areas (Supplementary Fig. 6d; regression slope: 0.0 ± 0.2, *p* = 0.99, *t* = −0.02; intercept difference with respect to V1: LI −0.02 ± 0.06, *p* = 0.7, *t* = - 0.36; TO −0.01 ± 0.06, *p* = 0.89, *t* = −0.15; df = 56), thus behaving very differently from what observed in our recordings (Fig. 4d) and in the Allen data (Fig. 7d and 8b, left).

Given that the movies tested in [59] were much shorter (5 s) than those used in our experiments (20 s) and in the Allen study (120 s), and fewer trials were collected (10) as compared to our recordings (30), we checked whether these results could be explained by a lack of statistical power in the head-fixed rat dataset to detect the trends that were found in our recordings and in the Allen data. To this aim, we replicated the analysis of our recordings shown in Fig. 4b, d, after subsampling the number of available trials and progressively reducing the duration of the responses included in the analysis (see Supplementary Information and Supplementary Fig. 7 for details). We found that lowering the movie duration (and, to a lesser extent, the trial number) strongly reduced the sensitivity to detect the dependence of the timescale of the response from that of the movie, as well as the increase of response stability across the visual hierarchy (see Supplementary Fig. 7a, b). Conversely, the sensitivity to detect the differences between the intrinsic temporal scales of the extrastriate lateral areas and those of V1 was only marginally reduced (see Supplementary Fig. 7d-f). This suggests that the lack of increase of the intrinsic timescales from V1 to LI/TO in the head-fixed rat dataset is a genuine difference with our recordings in anesthetized rats and with the Allen experiments in awake, behaving mice – a difference that is fully consistent with the much-reduced increase of the intrinsic temporal scales observed in the Allen data when mice deliberately chose to rest rather than run (see Fig. 8b).

## Discussion

Most popular models of the ventral stream are purely feedforward and view the sensory hierarchy as implementing bottom-up computations that are driven primarily by the visual input and that extract representations that are increasingly invariant to image transformations [1; 5; 9; 19]. Most of these models are state-independent (or “static”) – once the model (typically a multilayer, feedforward neural network) has been trained, the instantaneous representation of the current visual input is independent from the previous activity of the units or the past history of the stimulus. This picture of the ventral stream suggests that responses in deeper areas will be more invariant to differences between static images (transformations) of the same object, as observed in several experiments in both monkeys and rats [6; 7; 10; 53; 54]. As a result, responses to dynamic movies should be more temporally stable in deeper areas. On the other hand, adaptive and top-down mechanisms (e.g., prediction error signals) have been identified in visual cortex [23; 69] that favor the encoding of surprising or transient inputs over predictable or sustained ones [27; 30; 37; 60; 70]. Unfortunately, the role played by these mechanisms in representing dynamic inputs is poorly studied, but an insight can be gained by running simulations of a recent model of neuronal adaptation [39], which we extended with the addition of neuronal noise and intrinsic integration (see Supplementary Information). The progressive shortening of the timescales we observed for the responses of the simulated units as a function of the strength of adaptation (Supplementary Fig. 8e) suggests that the latter could indeed enforce a common, similarly fast stimulus-driven dynamics throughout visual cortex.

In our study, we tested which of the above scenarios is more consistent with the dynamics of neuronal representations recorded along the rodent analogue of the ventral stream, while anesthetized rats were subjected to dynamic visual stimulation with naturalistic and synthetic movies. We found that, in all tested visual areas, the temporal scale of visual encoding depended on the characteristic timescale of the dynamic stimuli, yet it significantly increased from V1 to extrastriate ventral cortex (Figs. 4a, b and 5b, c). Such an increase along the hierarchy was particularly strong when we tested the ability of the cortical representations to sustain the discrimination of movie segments of similar duration that contained single objects translating along either similar or opposite trajectories within the visual field (Fig. 6b, c). This implies that the increase of invariance afforded by feedforward computations along the rat ventral pathway is not completely washed out by adaptative, recurrent and top-down processes. As a result, when the pathway is probed with dynamic stimuli, a functional hierarchy of representational timescales is observed that parallels the hierarchical buildup of invariance revealed by brief presentation of static stimuli [53; 54].

As an alternative explanation, one could wonder if this trend simply reflects a systematic increase of the width of the temporal filters that describes the temporal integration of low-level visual features by neurons along the hierarchy. In the Allen dataset, [60] estimated the spike autocorrelation functions of the recorded neurons during responses to flash stimuli, from which they extracted a characteristic time constant by fitting a decaying exponential. They found the resulting time constants to increase along the visual hierarchy. This raises the question of whether this increase can explain the growth of the response timescale to the movies, given that the two temporal scales were significantly correlated (see Supplementary Fig. 9a; *r* = 0.75; *p* < 0.01, 1-tailed t-test) even though they do not necessarily measure the same quantity. To verify this, we modeled each recorded neuron in the Allen dataset as a linear filter, with the spatial kernel given by its RF profile and increasingly broad exponential temporal kernels defined by the time constants estimated in [60]. By convolving the input signal (i.e., the 120 s-long movie) with these spatiotemporal linear filters, we obtained a crude simulation of the responses of the units to the movie. The resulting response timescales, estimated in the same way as in our other analyses, did not differ much among the areas. As a result, when we plotted them as a function of the anatomical hierarchy score, we did not observe any monotonic increase (Supplementary Fig. 9b, c). This suggests that the temporal filtering properties of rodent visual neurons do not play a key role in leading to the hierarchical increase of response timescales under dynamical stimulation.

A second key question addressed by our study is the role of those activity-dependent processes that can extend the persistence of neuronal representations along an object-processing pathway, affecting the temporal span over which fluctuations of the firing rate are correlated within a single trial. In both primates and rodents, the temporal scales of these intrinsic fluctuations have been found to increase along various cortical hierarchies [40–43]. Mechanistically, the nature of these intrinsic processes has not been clarified yet, but most authors attribute them to temporally extended input integration or to recurrent computations, meant to sustain the neuronal representation over a time scale that is suitable to guide perception and behavior [42; 43; 64]. Not surprisingly, this topic has received a lot of attention in the computational neuroscience literature. In fact, deep convolutional neural networks have emerged as successful models of the ventral stream [9], and authors investigating the limitations of purely feedforward architectures within this family have proposed including temporal dynamics and adaptive mechanisms [39], or recurrent computations [71–75]. Experimentally, however, most work has focused on static stimuli, and thus it is not known whether intrinsic dynamics contribute to the structure of population codes for stimuli that are themselves varying in time.

Our experiments addressed this question. We found strong evidence that intrinsic timescales increase along rat lateral extrastriate areas (Fig. 4c, d), and that such increase plays an incrementally important role in stabilizing the neural representation of the visual input along the cortical hierarchy (Fig. 5d and 6d). Also, we found a weak but significant dependence of the intrinsic timescales from the dynamics of the movies (Fig. 4d) – a trend that was also consistent with the magnitude of the intrinsic timescales measured in awake mice (see the next Section and Fig. 7f). This means that the intrinsic fluctuations of neuronal firing are correlated over a time span that is related to the overall rate of change of the visual input. Thus, although our experiments did not investigate the mechanisms behind such intrinsic, activity-dependent processes, they clarify their possible functional role and their relationship with the overall dynamics of the visual input.

To achieve precise control and repeatability of the stimuli falling within each neuron’s receptive field over an extended period (1.5/2 hours) we performed our acute recordings under anesthesia. We expected, in view of previous work in anesthetized animals [47; 62], that the spatiotemporal characteristics of population activity that we measured would be similar in awake animals. Indeed, our estimates of single-cell intrinsic processing timescales fall squarely within the ranges reported for behaving mice and monkeys [42; 43; 60]. On the other hand, our preparation may have suppressed top-down effects that could appear or be stronger in awake animals. Such effects would add to the intrinsic processing, whose importance, relative to the feedforward drive, increases in higher areas of the ventral stream, as already discussed.

On the basis of these considerations, we repeated our assessment of the temporal stability of neuronal firing during presentation of natural movies using the Allen dataset [60]. These data allowed testing the generality of our conclusions in a different rodent species (the mouse) in the awake state and along a visual hierarchy that is very well established at the anatomical level. In addition, by partitioning the recorded neuronal responses into resting and running epochs, we could compare the dynamics of neuronal firing during active and quiet wakefulness. We found the increase of timescales to be preserved along the mouse visual system, both in terms of stimulus-driven responses and intrinsic fluctuations (Fig. 7c, d). The hierarchical increase of the response timescales was actually more evident of that observed along rat lateral extrastriate areas. This is likely due to the fact that the Allen dataset includes also LGN (a very low-level, subcortical region) and encompasses a larger number of visual cortical areas, thus allowing the temporal scale of neuronal responses to be tested across a deeper processing hierarchy. We also found the increase of temporal scales to be much attenuated (especially in the case of the intrinsic dynamics) during bouts of quiet wakefulness, as compared to epochs of active wakefulness (Fig. 8). To further investigate the state dependency of our findings, we analyzed a third dataset of recordings performed in awake but body restrained rats [59], finding no evidence of an increase of response and intrinsic timescales from V1 to extrastriate lateral areas (Supplementary Fig. 6). Although the failure to detect a growth of the response timescales could in part be explained by a lack of statistical power (Supplementary Fig. 7), this argument does not apply to the intrinsic timescales. Therefore, the fact that they remained stable across the lateral processing hierarchy appears to be a genuine result of the enforced immobilized state – a finding that is fully compatible with the strong attenuation of the hierarchical growth of the intrinsic timescales observed in the Allen dataset during the resting bouts (Fig. 8b).

Importantly, these results fit with those of previous studies reporting weakened processing of the visual input by rodent visual neurons (e.g., in terms of response magnitude, spatial integration, sparseness, reproducibility and discriminatory power) during quiet wakefulness, as compared not only to active wakefulness, but also to the anesthetized state [44; 46; 47]. These findings led some authors to conclude that during quiet wakefulness visual cortex may be perceptually disengaged or detached from the visual environment, even relative to the anesthetized state [44; 47]. The reduced (Fig. 8b) or canceled (Supplementary Fig. 6c, d) hierarchical growth of the intrinsic timescales observed in our analyses during, respectively, voluntary or enforced stillness, is highly consistent with this interpretation. In fact, during quiet wakefulness, there is arguably no need to regulate the temporal span over which the incoming visual signal gets integrated to guide perceptual decisions and motor behavior, especially in the case of enforced body restraint. As such, we could speculate that it could be computationally and metabolically efficient for the brain to turn off those activity-dependent, intrinsic processes that set the proper timescales of signal integration across the various stages of the visual processing hierarchy.

But what roles specifically could these intrinsic dynamics play in behavior? Intrinsic processing may be crucial for perception in a noisy, changing environment. For example, maintenance of sensory information by stimulus-independent temporal correlations in population activity can lead to better behavioral performance, when consistent estimates of a quantity are needed [43; 64]. Alternatively, intrinsic processing may support predictive coding [36; 38], allowing neural circuits to use feedforward inputs to predict and represent what will happen next, an ability with obvious utility for behavior. In order to play these roles effectively, the dynamics of intrinsic processing should be adapted to the temporal structures that are typically encountered in natural environments. Previous work has shown (or at least suggested) that many aspects of spatial and temporal processing in the early visual system are indeed adapted to the structure of natural scenes. Examples are histogram equalization [76] and On-Off circuit asymmetries in the retina [77], development of V1 simple and complex cells [57; 78–81], texture perception [82], as well as eye movements [83]. The idea that intrinsic processing could be similarly adapted deep into the cortical hierarchy could be causally tested by altering the animals’ visual environment during development or while learning a task, and then measuring neural population activity across the cortical hierarchy with ethologically relevant, dynamic stimuli.

## Methods

### Animal preparation and surgery

All animal procedures were in agreement with international and institutional standards for the care and use of animals in research and were approved by the Italian Ministry of Health: project N. DGSAF 22791-A, submitted on Sep. 7, 2015 and approved on Dec. 10, 2015 (approval N. 1254/ 2015-PR). 18 male Long Evans rats (Charles River Laboratories) with age 3-12 months and weight 300-700 grams were anesthetized with an intraperitoneal (IP) injection of a solution of 0.3 mg/kg of fentanyl (Fentanest®, Pfizer) and 0.3 mg/kg of medetomidine (Domitor®, Orion Pharma). Body temperature was kept constant at ~37° with a warming pad, and a constant flow of oxygen was delivered to the rat to prevent hypoxia. The level of anesthesia was monitored by checking the absence of tail, ear and hind paw reflex, as well as monitoring blood oxygenation, heart and respiratory rate through a pulse oximeter (Pulsesense-VET, Nonin), whose sensor was attached to one of the hind paws. After induction, the rat was placed in a stereotaxic apparatus (Narishige, SR-5R) in flat-skull orientation (i.e., with the surface of the skull parallel to the base of the stereotax), and, following a scalp incision over the left and posterior parietal bones, a craniotomy was performed over the target area in the left hemisphere (typically, a 2×2 mm2 window). Dura was also removed to ease the insertion of the electrode array. Stereotaxic coordinates for V1 recordings ranged from −7.49 to −8.36 mm anteroposterior (AP), with reference to bregma; for extrastriate areas (LM, LI and LL), they ranged from −6.64 to −7.74 mm AP.

Once the surgical procedure was completed, the stereotaxic apparatus was moved on top of an elevated rotating platform. The right eye was immobilized with an eye ring anchored to the stereotaxic apparatus, and the left one was covered with opaque tape. The platform was then rotated, so as to align the right eye with the center of the stimulus display and bring the binocular portion of its visual field to cover the left side of the display. Throughout the experiment, ophthalmic solution Epigel (Ceva Vetem) was regularly applied onto the right eye and the exposed brain surface was covered with saline to prevent drying. In addition, a constant level of anesthesia was maintained through continuous IP infusion of the same anesthetic solution used for induction, but at a lower concentration (0.1 mg/kg/h fentanyl and 0.1 g/kg/h medetomidine), by means of a syringe pump (NE-500; New Era Pump Systems).

### Neuronal recordings

Extracellular recordings were performed with 32-channel silicon probes (Neuronexus Technologies), following the same procedure used in [53]. Briefly, to maximize the receptive field coverage, recordings in V1 were performed with 4- or 8-shanks probes, which were inserted perpendicularly into the cortex. Recordings from lateral extrastriate areas were performed using single-shank probes, which were inserted diagonally into the cortex with an angle of ~30°, in order to map the progression of receptive fields’s centers along the probe and track the reversal of retinotopy between adjacent areas (see [53; 55]). The space between recording sites on each shank ranged from 25 to 200μm; the distance between shanks (when more than one) was 200μm; the surface of recording sites was either 177 or 413 μm^2^. Extracellular signal was acquired with a TDT system 3 workstation (Tucker-Davis Technologies) at a sampling rate of ~24kHz. Before insertion, the probe was coated with Vybrant® DiI cell-labeling solution (Invitrogen, Oregon, USA), to allow visualizing the probe insertion track *post-mortem*, through histological procedures.

A total of 19 rats were used for this study. Since, in some occasions, multiple recording sessions were performed from the same animal, this yielded a total of 23 sessions. After spike sorting (see below) these recordings yielded a total of 1313 well-isolated single units: V1 (510), LM (126) LI (209) LL (401). Following the application of a reproducibility filter on the neuronal responses of the units across repeated presentations of the movie stimuli (see below), the final count of single units that were further analyzed in our study was 294, of which 168 in V1 (5 different rats, for a total of 7 recording sessions), 20 in LM (4 rats, 4 sessions), 36 in LI (8 rats, 8 sessions), 70 in LL (11 rats, 12 sessions).

### Visual stimuli

Stimuli were presented full-field to the right eye at a distance of 30 cm on a 47-inch LCD monitor (SHARP PN-E471R), with 1920 × 1080 resolution, 60Hz refresh rate, 9ms response time, 700cd/m2 maximum brightness, 1200:1 contrast ratio, spanning a field of view of ~120° azimuth and ~89° elevation. The stimuli were presented using Psychtoolbox [84] in MATLAB in a pseudorandom order. Between each stimulus (movie or drifting bar) a black screen was shown for at least 200 ms.

The receptive fields of the neurons recorded along a probe were estimated by showing drifting oriented bars over a grid of 73 locations on the stimulus display, covering the central 110° azimuth span and central-upper 70° elevation span of the total field of view of the monitor. The bars were 10º long and drifted 7 times along four different directions (0°, 45°, 90°, and 135°). A spherical correction (as described in [85]) was applied to each bar to compensate for shape distortions at large eccentricities. As shown in Fig. 2, mapping the inversion of the retinotopic map at each area boundary allowed identifying the visual area each neuron was recorded from [53–55].

The main stimulus set consisted of 9 movies: two fast and two slow *manual* movies, two *ratcam* movies, a phase-scrambled version of one of the fast movies and a phase-scrambled version of one of the ratcam movies (see next paragraph for details). The resolution of these movies was 720 × 1280 pixels and they were presented for 20 seconds at a rate of 30 frames per second (fps). An additional white-noise movie with a resolution of 45 × 80 pixels was shown at 20 fps for 20 seconds. All movies were converted to grayscale and were gamma-corrected offline with a lookup table calculated for the monitor used for stimulus presentation. During the course of the experiment, each movie was presented 30 times. The videos can be found in the supplementary material.

The manual movies were designed to reproduce the continuous flow of visual information typically experienced by an observer that explores an environment containing a number of distinct visual objects, placed in different locations. To this aim, we 3D-printed various geometrical objects, we painted them black or white (see examples in Fig. 3a), we placed them inside an arena and we simulated an observer roaming through such an environment by smoothly moving a hand-held camera across the arena for 20 s. The camera was moved at two different speeds to obtain slower and faster movies. The *ratcam* movies were intended to better simulate the visual input of a rat exploring a natural environment [47]. They were obtained by placing a small web camera (Microsoft Lifecam Cinema HD) on the head of a rat, while it was running freely inside the arena that contained some of the 3D-printed objects and another rat.

The phase-scrambled movies were obtained by performing a spatiotemporal fast Fourier transform (FFT) over the standardized 3D array of pixel values, obtained by stacking the consecutive frames of a movie. The phase spectrum was extracted and shuffled, then merged with the unaltered amplitude in such a way to preserve the conjugate symmetry of the initial transform. An inverse FFT was then applied on the resulting array. The imaginary parts were discarded, then the original mean and standard deviation were restored. Values outside the 0-255 range were clipped in order to obtain a valid movie. The white noise movie was generated by randomly setting each pixel in the image plane either to white or black.

### Histology

In a few experiments (5 rats), at the end of the recording session, each animal was deeply anesthetized with an overdose of urethane (1.5 gr/kg) and perfused transcardially with phosphate buffer saline (PBS) 0.1 M, followed by 4% paraformaldehyde (PFA) in PBS 0.1 M, pH 7.4. The brain was then removed from the skull, post-fixed in 4% PFA for 24 h at 4°C, and then immersed in cryoprotectant solution (15% w/v sucrose in PBS 0.1 M, then 30% w/v sucrose in PBS 0.1 M) for 48 h at 4 °C. The brain was finally sectioned into 30μm-thick coronal slices using a freezing microtome (Leica SM2000R, Nussloch, Germany). Sections were mounted immediately on Superfrost Plus slides and let dry at room temperature overnight. A brief wash in distilled water was performed, to remove the excess of crystal salt sedimented on the slices, before inspecting them at the microscope. Each slice was then photographed with a digital camera adapted to a Leica microscope (Leica DM6000B-CTR6000, Nussloch, Germany), acquiring both a DiI fluorescence image (700 nm DiI filter) and a bright-field image at 2.5X and 10X magnification. By superimposing the fluorescence and bright-field images, we reconstructed the tilt and the anteroposterior (AP) position of the probe during the recording session (se example in Fig. 2a, left).

### Data analysis

#### Spike sorting and selection of the units included in the analyses

Data were spike-sorted offline with the spike sorting package KlustaKwik-Phy [86] in two steps: automated spike detection, feature extraction and expectation maximization (EM) clustering were followed by manual refinement of the sorting using a customized version of the GUI. The last step was performed by taking into account many features of the candidate clusters: 1) the distance between their centroids and their compactness in the space of the principal components of the waveforms (a key measure of goodness of spike isolation); 2) the shape of the auto- and cross-correlograms (important to decide whether to merge two clusters or not); 3) the variation, over time, of the principal component coefficients of the waveform (important to detect and take into account possible electrode drifts); and 4) the shape of the average waveform (to exclude, as artifacts, clearly non-physiological signals). Clusters suspected to contain a mixture of one or more units were separated using the “reclustering” feature of the GUI (i.e., by rerunning the EM clustering algorithm on the spikes of these clusters only).

At the end of the manual refinement step, we further screened the resulting well-isolated single units to make sure that their firing was reproducible across repeated stimulus presentations. The selection was based on a reproducibility index that quantified, for each neuron, how reliable its responses were to its preferred frames within a movie (see definition below). This metric was chosen over other approaches that rely on computing the correlation coefficient between responses to repetitions of the same stimulus [59; 87], because we wanted to avoid the inclusion of silent neurons that would have yielded high correlations due to lack of activity. To calculate the reproducibility index for a given unit and movie, we included in the metric the top 10% time bins with the highest neuronal response (given by the firing rate of the neuron in the bins). Let 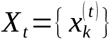 be the set of responses of the neuron 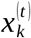 in one of such preferred time bins *t* (with *k* denoting the stimulus repetition). Let *N* be the number of repetitions. Then, the reproducibility index is calculated as 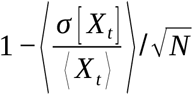. The resulting metric ranges from 0 to 1 [88], where 1 corresponds to ideal responses with perfectly reproducible trials. A cell was considered reproducible if, for at least one of the nine movies, the reproducibility index was 0.7. This threshold was arbitrarily set following an extensive visual inspection of raster plots from different movies to ensure that clearly responsive and reproducible cells were included. For all the analyses described in this study, the responses of these units during the presentation of the movie stimuli were converted to a spike-count representation by binning spike trains in 33ms bins.

#### Characteristic timescale of the movie stimuli

To quantify how fast the pixel-level content of the movies changed over time, we computed the following metric. Given a movie, for a given time lag *Δ*, we computed the average correlation coefficient between all movie frame pairs separated by that lag, on a pixel-wise basis, i.e.:

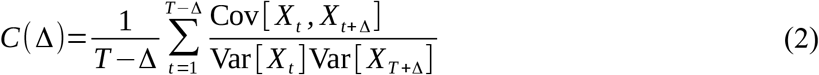

where *T* is the total number of frames in the movie, 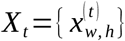 is the movie frame *t* and expectations are taken over pixel positions, indexed by *w,h* (for width and height). We then plotted the correlation *C* (*Δ*) as a function of *Δ* (Fig. 3b). To fit the decay of the correlation with *Δ* we considered two possible functional forms: a decaying exponential of the form *y* (*t*)=*a* exp (*−t* / *τ*) and a damped oscillation function of the form *y* (*t*)=*a* exp (*−t* / *τ*) cos (*ωt* + *ϕ*). Fitting was performed with the basin-hopping algorithm [89] coupled with L-BFGS-B [90]. Model selection was performed independently for each fit with an Extra Sum of Squares test [91], choosing the more complex model whenever the p value from the test resulted below the threshold of 0.05. The model selection procedure selected the simple exponential form in all cases. The time constant *τ* of the exponential decay was taken as the characteristic timescale of the stimulus.

#### Characteristic timescale of stimulus-driven neuronal population responses

To measure the characteristic timescale of neural stimulus correlations from population peri-stimulus time histograms (PSTHs) in our anesthetized rat data, we applied a very similar procedure to that described above for the visual input. We used the same expression as above (Eq. 2) to compute *C* (*Δ*), but now we set 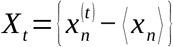, where 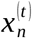 is the trial-averaged spike count of cell number *n* at time frame *t*, and subtracting ⟨ *x*_n_ ⟩ centers 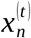 around its temporal average 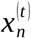 as a function of *t* is color-coded in Fig. 3c for the V1 and LL populations, in response to two example movies). A time constant was extracted by fitting either an exponential decay of the form *y* (*t*)=*a* exp (*−t* / *τ*) or a damped oscillation function of the form *y* (*t*)=*a* exp (*−t* / *τ*) cos (*ωt* + *ϕ*)+ *b* for *t* >0 (see Fig. 4a). Model selection was carried out as described in the previous section. The time constant *τ* of the exponential decay or of the exponential envelope of the damped oscillation was taken as the characteristic timescale of the trial-averaged population response (PSTH). The simple exponential form was selected for three of the movies in V1 and one of the movies in LL; the damped oscillation form was selected for all other movies. We note that here, as well as in the case of intrinsic time scale estimation (see below), the model selection procedure ensured that the estimated timescale matched the characteristic time over which the statistical dependence between the signal *X*_*t*_ at time *t* and the signal *X*_*t*_ + τ at time *t+τ* became small compared to its value at short lags. Crucially, the model selection procedure allowed this to hold even in presence of oscillatory patterns in the correlation functions, where the fact that the correlation goes to zero at a certain lag can hide the fact that it takes on significant negative values (indicating the persistence of a statistical relationship, albeit with flipped sign) and can further possibly come back to positive values at larger lags. We considered one population PSTH for each cortical area we recorded from, pooling together all the available sessions, as illustrated in Fig. 3c.

#### Intrinsic timescale of neuronal activity

Intrinsic timescales of neuronal activity were computed in a way that mirrors the stimulus-driven correlation timescales defined in the previous section (see Supplementary Fig. 1) and that is a simple extension of the definition given by [42]. To compute *C* (*Δ*) for a given cell, we set 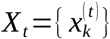 in Eq. 1, where 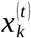 is the spike count of the cell at time bin *t* in trial *k*. Following [46], we then averaged *C* (*Δ*) across all units belonging to the same area before extracting a time constant (see example in Fig. 4c). The characteristic timescale of intrinsic correlation decay was computed as above for the PSTH, using the same fitting functions and the same model selection procedures. Intrinsic correlations in V1 were well fitted by simple exponential decays, while the damped exponential form was selected for fitting intrinsic correlations for all movies in extrastriate areas.

#### Characteristic timescale of decoding performance

After computing the average classifier performance *p* (*Δ*) as a function of the lag *Δ* (see next section), a characteristic decay time constant was extracted by the same procedure previously used to compute the timescales of the movies, of the stimulus-driven responses and of the intrinsic activity. Namely, either an exponential decay of the form *y* (*t*)=*a* exp (*−t* / *τ*)+*b* or a damped oscillation function of the form *y* (*t*)=*a* exp (*−t* / *τ*) cos (*ωt* + *ϕ*)+ *b* were fitted to *p* (*Δ*) for *Δ* >0 (see example in Fig. 5b). Model selection was performed as described above, and the exponential decay time constant *τ* was extracted as the quantity of interest. The majority of pseudopopulations/movie/trial set combinations were fit with the damped-oscillatory functional form (V1: 108 out of 144; LM: 14 out of 18; LI: 17 out of 18; LL: 48 out of 54).

#### Classifier analysis

To assess the discriminability of population activity at different points in time during the presentation of a given movie, we proceeded as follows. For each recording session, we divided the 30 available recording trials in a training set of 20 trials and a validation set of 10 trials. We pooled all units that were recorded from a given area across all sessions to obtain a total number of units *N* per area. We generated K pseudopopulations of *M* < *N* units (with *M* = 20 for most of the analyses) by selecting as many random non-overlapping subpopulations of size M as possible. For instance, in V1 K=8 as N=168. These subpopulations were the same in the training and in the validation set. Following standard procedure when working with pseudopopulations, for each trial set (training and validation) we shuffled cell activity across trials to destroy cross-cell noise correlations.

We then considered all pairs of time bins along a trial that were separated by at least 40 time bins (i.e., by 1320 ms, as bin size was 33 ms). For each of these “reference” pairs of time bins (gray boxes in Fig. 5a), we trained a linear support vector classifier (provided by liblinear [92] via scikit-learn [93]) to discriminate population activity samples from one of the element of the pair vs. the other. The penalty hyperparameter was chosen by 3-fold cross-validation within the training set, performing a grid search over candidate values 10^-2,-1,0,1,2^. After selecting the best value of the hyperparameter, the classifier was re-trained on the full training set.

To assess the temporal stability of the population activity, the trained classifier was then tested on samples of population activity coming from different time bins than those it was trained on (orange and green boxes in Fig. 3A). More specifically, if a classifier was trained on the pair of time bins at times t_1_ and t_2_, it was then tested on pairs of time point at times t_1_+*Δ* and t_2_+*Δ*, where *Δ* took on values {−20,…,−1,0,1,…,20} bins * 33 ms/bin. The fraction of correctly decoded trials at negative and positive time increments was then averaged to yield a classifier performance curve p(*Δ*) with *Δ* ranging from 0 to 20 bins*33 ms/bin = 660 ms. This performance curve was computed for all possible reference pairs of time bins, and averaged over all pairs. The resulting average performance decay curve (for each individual movie stimulus and pseudopopulation) was used to compute the typical timescale of self-similarity for population activity as described above. Examples of such performance decay curves are given in Fig. 5b for the case in which the test was performed on held-out trials (i.e., green boxes in Fig. 5a).

This analysis was performed separately on the training and the held-out trial sets (respectively orange and green boxes in Fig. 5a), while keeping the trained decoders fixed (see performance decay curves in Fig. 5b and Supplementary Fig. 3a). When computed on the training set, p(0) corresponds to the performance of the classifier on its own training data, but p(33ms) is the performance of the classifier on an entirely new set of data, which happens to come from the same set of experimental trials as the training data. When compared to the results on the held-out set, this allowed quantifying the extent to which the self-similarity of the population activity in time was due to the stimulus-locked representational structure of the population code rather than to intrinsic, within-trial, temporal correlations in the activity of each neuron. Intuitively, population activity at time *t* in trial *i* can be expected to be more similar to population activity at time *t+Δ* in the same trial than in a different trial, precisely because of the existence of within-trial, stimulus-independent correlations in the activity of each neuron. Comparing the performance of the classifier on the training set vs. the held-out validation set, as done in Fig. 5d, allowed therefore to assess the importance of this correlational structure in enhancing the self-similarity of population activity over time (see Eq. 1).

#### Regression analysis and hypothesis testing

Linear regression for the dependence of the timescale of stimulus-driven responses and intrinsic activity on the stimulus timescale (Fig. 4,b, d), as well as for the dependence of the amount of classifier performance due to intrinsic correlations on the timescale of intrinsic activity (Fig. 5d and Supplementary Fig. 2c) was computed by ordinary least squares. In each case, residuals were not incompatible with a normal distribution (Jarque-Bera, p > 0.05). Regression and tests were performed with the Python package *statsmodels*.

The distribution of the data was less regular for the case of the classifier performance vs. the timescale of the movie (Fig. 5c and Supplementary Fig. 3b). In particular, there were a few outliers with very long decoding timescales, especially for LL, which could have biased our conclusion in favor of the hypothesis that higher areas in the hierarchy have longer processing timescales. To prevent our conclusions from relying disproportionately on these points, we performed the corresponding linear regressions using a robust estimator (Theil-Sen estimator; [94]). The confidence intervals were determined by percentile bootstrap (10000 samples; [95]) stratified by cortical area. This combination of estimator and CI determination procedure has been shown to provide reasonable coverage probability in the face of model misspecification and data contamination by outliers [94]. Following [95], p-values were computed by subtracting the bootstrap distribution of any given parameter from its estimated value to obtain a surrogate for the null distribution. The p-value was then obtained as the mass of the surrogate distribution for values larger than the estimate. This yielded a one-tailed test where the null hypothesis was that the parameter value was not larger than zero. Regression and tests were performed with custom R code [96] using the *boot* and *WRS* libraries.

#### Classifier analysis on hand-matched movie segments

The results illustrated in Fig. 6 were obtained by performing a variant of the classifier analysis on a small subset of neuronal recordings, corresponding to the presentation of movie segments that were carefully matched in pairs to reproduce as much as possible the conditions under which representation invariance is typically studied with static stimuli. All movie segments were taken to be exactly 29 frames (~967ms) long, and as described in the main text they were all selected to show a single object, moving smoothly from the left to the right of the frame (or vice versa) over the course of the segment. The object could be black or white, leading to the existence of four categories of movie segment pairs, based on the match or mismatch between the color of the object in each element of the pair and their direction of motion: (1) opposite color, same direction; (2) opposite color, opposite direction; (3) same color, same direction; and (4) same color, opposite direction. The number of available movie segment pairs was 4, 6, 5 and 6 respectively for the four conditions. For each segment pair and each cortical area, a classifier analysis similar to the one outlined above was performed, as follows. First of all, as above, K non-overlapping neural pseudopopulations of 20 cells each were formed at random, where K=floor(N/20) and N was the total number of available cells (for area LM, this just meant forming one pseudopopulation taking up all available cells). Neural activity was binned in temporal bins of duration 1/30s, (rather than 33ms as in the other analyses), corresponding to the exact duration of each movie frame, to ensure straightforward alignment of neural binned data and movie frames. Each of the time bins spanning the duration of either movie segment was sequentially considered as the training bin for the classifier, which was then tested on the other bins, yielding a performance vs lag curve, where the lag could go from −28 bins to +28 bins depending on the identity of the training bin. For instance, when the classifier was trained on the first bin of both segments, the classifier’s performance would be evaluated for lags spanning the range from 0 to +28. On the other hand, when the classifier was trained on the middle bin of both segments (bin number 15), the performance vs lag curve would be computed for lags spanning from −14 to +14. All performance curves were then aligned and averaged (ignoring missing values), yielding a summary performance curve for lags spanning from - 28 to +28. This performance curve was further “folded” by identifying positive and negative lags. The folded performance curve (spanning now from 0 to +14 bins, i.e., from 0 to ~467ms) was then averaged across all neural pseudopopulations for the areas. Finally, this procedure was repeated 10 times, each time selecting different random pseudopopulation, and the resulting curves were averaged. This yielded an average performance curve for each movie segment pair and each brain area. For each of these curves, a characteristic timescale was computed using the same curve-fitting methods described in the previous section. A final estimate of the timescale for each condition was obtained as the average timescale derived from all movie segment pairs in that condition. Statistical significance of the difference between the timescales in V1 and those of the other areas was assessed by performing a simple linear regression of the timescales of the individual movie pairs versus area identity and condition. Mathematically, this was expressed as follows:

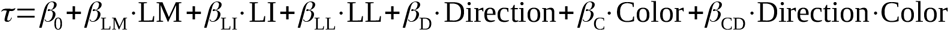

In this expression, LM (resp. LI, LL) is an indicator variable that is 1 when the area is LM (resp LI, LL) and 0 otherwise, and Direction (resp. Color) is an indicator variable that is 1 when the movement direction (resp. object color) is the same in the two movie segments making up a pair. Therefore, for instance, β_LM_ represents the mean difference in timescale between LM and V1 controlling for the effect of the condition (direction and color).

#### Analysis of awake mouse data

The awake mouse dataset (here and elsewhere referred to the Allen dataset) was collected and published by the Allen Institute for Brain Science. We refer to the original publication [60] for an exhaustive description of the experimental procedures and data processing.

The Allen dataset is subdivided into individual sessions. We used the Allen Software Development Kit (Allen SDK) to extract stimulus-aligned spike timings for the relevant visual areas (LGN, V1, LM, RL, AL, PM and AM) from the first 17 sessions included in the dataset and we pooled all units within each area. Firing rates were binned using a bin size of 33ms, consistent with our other analyses. As detailed in the Results section, we only considered the neural activity recorded in response to the presentation of one of the two movies in the dataset (the longest of the two, named Natural Movie 3), as the other movie (named Natural Movie 1) was composed by highly static shot, leading to an irregular structure of the temporal pixel correlations (Supplementary Fig. 4a, left) that when collapsed as a correlation function were not well described by any of the functional forms we considered (decaying exponential, damped oscillation; see Supplementary Fig. 4b, red curves). We analyzed the movie frames and the neural data using the same methods detailed above for the anesthetized rat experiment, with a few differences. First, as we only had access to one movie, we did not study the dependence of the timescales on the stimulus timescale, but we only analyzed the difference in timescales across areas. Second, all correlation functions (pixel correlations in the stimulus, as well as response and intrinsic correlations in the neural data) were well fit by simple decaying exponentials, so the damped oscillation functional form was never used.

To study the variation of the neural timescales along the visual pathway, we performed an ordinary least squares linear regression of the timescales against the anatomical score of their respective area.

#### Comparison of running and resting conditions in awake mouse

Given a recording session, we used the velocity information v to identify epochs of the neuronal responses to the movies during which the mouse was either resting (v < 1 cm/s) or running (v > 1 cm/s). Each session had a number of these epochs, and by construction each rest epoch divided two adjoining running epochs. In addition, given that each movie was presented 10 times within a session, the segments of the movie in which the mouse was at rest or running were different in each presentation. Since our analysis required processing the responses of each recorded neuron across multiple repeated presentations (or trials) of the same stimuli, we looked for segments of the movie where the resting (or running) epochs obtained for the various trials overlapped and we took their intersection (see Supplmentary Fig. 5). The number of resting (or running) epochs shared across trials decreased as a function of the number of trials considered and, concomitantly, their duration became shorter. In principle, our comparison between the running and the resting conditions could have been affected by our specific choice in this trade-off between number of trials and amount of available data (duration and number of the shared resting or running epochs). To avoid any such bias, we repeated our analysis for any combination of four choices of trial number *N*_*t*_ (i.e., 3, 4, 5 and 6 trials) and five choices of the minimal duration L of the epochs to be included in the analysis (i.e., 2.5, 3.3, 4.1, 5.0 and 6.6 s). For each of these combinations, we computed the stimulus-driven and intrinsic correlation functions, we fitted the exponential decays and we obtained their time constants. This procedure was repeated for all the 17 recording sessions of the Allen dataset included in our analysis, so as to obtain an estimate of the average temporal stability of neuronal activity across the 7 visual areas during quiet wakefulness (i.e., during the resting epochs) as well as during active wakefulness (i.e., during the running epochs). To make sure that the time constants included in the final averages were obtained from exponential fits with similar quality, for any combination of the parameters *N*_*t*_ and L we pooled the errors of the fits obtained across all the resting and running epochs of all areas. We then looked at the resulting, overall distribution of fit errors and only retained the time constants of those fits with an error in the lower 50th percentile. Since in general, for any given area, the numbers of resting and running epochs yielded by this selection procedure were different, we equated them by subsampling the epochs of the more populate state. This ensured that the number of time constants used to assess the neuronal dynamics of a given area was the same for the resting and running state.

#### Analysis of awake rat data

For the awake rat analyses of Supplementary Fig. 6, we used the neuronal recordings collected in [59], for which we refer for a detailed description of the acquisition methodology. In brief, single-unit recordings were performed from areas V1, LI and TO on head-fixed, awake rats, while the animals were shown natural movie stimuli. The stimulus set included 20 movies, each 5 seconds long, and the number of recorded neurons was 50 in V1, 53 in LI, and 52 in TO. As for the anesthetized data, in our analyses the spike data was discretized in temporal bins 33ms long. We discarded all neurons that didn’t meet the reproducibility criterion described above (under *Spike sorting and selection of the units included in the analyses*), and this reduced the number of available neurons to 44 in V1, 40 in LI, and 38 in TO. Since the structure of the data was very similar to our new recordings in anesthetized rat, we were then able to apply the same analysis pipeline to estimate stimulus timescales, response timescales, and intrinsic timescales, leading to the data reported in Supplementary Fig. 6. Three exceptions were made to this general principle: 1) since the movies were shorter, we considered a smaller range of lags over which to compute the correlations (0.5s instead of 2s, compare Fig. 4a,c with Supplementary Fig. 6 a,c); 2) because of the particular trend typically exhibited by intrinsic correlations (with a peak at some nonzero but short lag, see Supplementary Fig. 6c), only the tail of the intrinsic correlations was considered, using only lags larger than 5 bins (=165ms); and 3) the form of the function used for fitting the correlations was now *y* (*t*)=*a* exp(−*t*/ *τ*)[*c* +cos (*ωt* + *ϕ*)]+*b*. All details of the regression analysis were the same as for the anesthetized data.

## Supporting information

Movie: manual fast 1

Movie: manual fast 2

Movie: manual slow 1

Movie: manual slow 2

Movie: ratcam 1

Movie: ratcam 2

Movie: white noise

Movie: phase-scrambled manual fast 1

Movie: phase-scrambled ratcam 2

## Data and code availability

The datasets and code generated during this study have been made available confidentially for peer review. They will be made public and permanently reachable upon publication of the paper. The Allen dataset is freely accessible online, as documented in [60]. The awake rat data from [59] can be found in [97] (doi:10.17605/OSF.IO/M2E6D).

## Acknowledgements

EP thanks Stefano Panzeri for many discussions, prior to this work, on temporally persistent neural codes and in particular on the topic of temporal noise correlations. We thank Walter Vanzella for his help in setting up the first version of the decoding analysis. We thank Margherita Riggi for her help in performing the histological procedures. This work was supported by: European Research Council (ERC) Consolidator Grant (DZ, project n. 616803-LEARN2SEE); NIH grant R01EY07977 (VB); NIH grant R01NS113241 (EP).

## Author Contributions

Conceptualization, E.P., L.S., V.B. and D.Z.; Data Curation, E.P., L.S, P.M. and K.V.; Formal Analysis, E.P., L.S., P.M. and R.C.; Funding Acquisition, E.P., V.B. and D.Z; Investigation, L.S.; Methodology, E.P., L.S., P.M., R.C., K.V., H.O.B., V.B and D.Z.; Project Administration, D.Z.; Resources, D.Z.; Software, E.P., L.S., P.M and R.C.; Supervision, D.Z.; Writing – Original Draft Preparation, E.P. and D.Z.; Writing – Review & Editing, E.P., L.S., P.M., K.V., H.O.B., V.B. and D.Z.

## References

[1] DiCarlo, J. J.; Zoccolan, D. and Rust, N. C. (2012). How does the brain solve visual object recognition? Neuron 73, 415–434.

[2] Zoccolan, D. (2015). Invariant visual object recognition and shape processing in rats. Behavioural brain research 285, 10–33.

[3] Leopold, D.; Mitchell, J. and Freiwald, W. (2020). Evolved Mechanisms of High-Level Visual Perception in Primates. In: Kaas, J. H. (Ed.), Evolutionary Neuroscience, Academic Press.

[4] DiCarlo, J. J. and Cox, D. D. (2007). Untangling invariant object recognition. Trends in cognitive sciences 11, 333–341.

[5] Riesenhuber, M. and Poggio, T. (1999). Hierarchical models of object recognition in cortex. Nature neuroscience 2, 1019–1025.

[6] Li, N.; Cox, D. D.; Zoccolan, D. and DiCarlo, J. J. (2009). What response properties do individual neurons need to underlie position and clutter “invariant” object recognition? Journal of neurophysiology 102, 360–376.

[7] Rust, N. C. and Dicarlo, J. J. (2010). Selectivity and tolerance (“invariance”) both increase as visual information propagates from cortical area V4 to IT. The Journal of neuroscience: the official journal of the Society for Neuroscience 30, 12978–12995.

[8] Pagan, M.; Urban, L. S.; Wohl, M. P. and Rust, N. C. (2013). Signals in inferotemporal and perirhinal cortex suggest an untangling of visual target information. Nature neuroscience 16, 1132–1139.

[9] Yamins, D. L. K.; Hong, H.; Cadieu, C. F.; Solomon, E. A.; Seibert, D. and DiCarlo, J. J. (2014). Performance-optimized hierarchical models predict neural responses in higher visual cortex. Proceedings of the National Academy of Sciences of the United States of America 111, 8619–8624.

[10] Hong, H.; Yamins, D. L. K.; Majaj, N. J. and DiCarlo, J. J. (2016). Explicit information for category-orthogonal object properties increases along the ventral stream. Nature neuroscience 19, 613–622.

[11] Berkes, P. and Wiskott, L. (2005). Slow feature analysis yields a rich repertoire of complex cell properties Journal of Vision 5, 9.

[12] Cadieu, C. F. and Olshausen, B. A. (2011). Learning Intermediate-Level Representations of Form and Motion from Natural Movies Neural Computation 24, 827–866.

[13] Einhäuser, W.; Kayser, C.; König, P. and Körding, K. P. (2002). Learning the invariance properties of complex cells from their responses to natural stimuli. The European journal of neuroscience 15, 475–486.

[14] Földiák, P. (1991). Learning Invariance from Transformation Sequences. Neural Computation 3, 194–200.

[15] Körding, K. P.; Kayser, C.; Einhäuser, W. and König, P. (2004). How Are Complex Cell Properties Adapted to the Statistics of Natural Stimuli? Journal of Neurophysiology 91, 206–212.

[16] Wallis, G. (1996). Using Spatio-temporal Correlations to Learn Invariant Object Recognition. Neural networks: the official journal of the International Neural Network Society 9, 1513–1519.

[17] Wallis, G. and Rolls, E. T. (1997). Invariant face and object recognition in the visual system. Progress in Neurobiology 51, 167–194.

[18] Wiskott, L. and Sejnowski, T. J. (2002). Slow Feature Analysis: Unsupervised Learning of Invariances Neural Computation 14, 715–770.

[19] Wyss, R.; König, P. and Verschure, P. F. M. J. (2006). A Model of the Ventral Visual System Based on Temporal Stability and Local Memory PLOS Biology 4, e120+.

[20] Krizhevsky, A.; Sutskever, I. and Hinton, G. E. (2017). ImageNet classification with deep convolutional neural networks. Communications of the ACM 60, 84–90.

[21] Grill-Spector, K.; Henson, R. and Martin, A. (2006). Repetition and the brain: neural models of stimulus-specific effects. Trends in cognitive sciences 10, 14–23.

[22] Kohn, A. (2007). Visual adaptation: physiology, mechanisms, and functional benefits. Journal of neurophysiology 97, 3155–3164.

[23] Webster, M. A. (2015). Visual Adaptation. Annual review of vision science 1, 547–567.

[24] Kaliukhovich, D. A.; De Baene, W. and Vogels, R. (2013). Effect of adaptation on object representation accuracy in macaque inferior temporal cortex. Journal of cognitive neuroscience 25, 777–789.

[25] Zhou, J.; Benson, N. C.; Kay, K. and Winawer, J. (2017). Unifying Temporal Phenomena in Human Visual Cortex. biorxiv preprint. doi:10.1101/108639

[26] Fritsche, M.; Lawrence, S. J. D. and de Lange, F. P. (2020). Temporal tuning of repetition suppression across the visual cortex. Journal of Neurophysiology 123, 224–233.

[27] Stigliani, A.; Jeska, B. and Grill-Spector, K. (2019). Differential sustained and transient temporal processing across visual streams. PLoS computational biology 15, e1007011.

[28] Stigliani, A.; Jeska, B. and Grill-Spector, K. (2017). Encoding model of temporal processing in human visual cortex. Proceedings of the National Academy of Sciences of the United States of America 114, E11047–E11056.

[29] Kaliukhovich, D. A. and Op de Beeck, H. (2018). Hierarchical stimulus processing in rodent primary and lateral visual cortex as assessed through neuronal selectivity and repetition suppression. Journal of neurophysiology 120, 926–941.

[30] Vinken, K.; Vogels, R. and Op de Beeck, H. (2017). Recent Visual Experience Shapes Visual Processing in Rats through Stimulus-Specific Adaptation and Response Enhancement. Current biology: CB 27, 914–919.

[31] Lueschow, A.; Miller, E. K. and Desimone, R. (1994). Inferior temporal mechanisms for invariant object recognition. Cerebral cortex (New York, N.Y.: 1991) 4, 523–531.

[32] Andrews, T. J. and Ewbank, M. P. (2004). Distinct representations for facial identity and changeable aspects of faces in the human temporal lobe. NeuroImage 23, 905–913.

[33] De Baene, W. and Vogels, R. (2010). Effects of adaptation on the stimulus selectivity of macaque inferior temporal spiking activity and local field potentials. Cerebral cortex (New York, N.Y.: 1991) 20, 2145–2165.

[34] Afraz, S.-R. and Cavanagh, P. (2008). Retinotopy of the face aftereffect. Vision research 48, 42–54.

[35] Afraz, A. and Cavanagh, P. (2009). The gender-specific face aftereffect is based in retinotopic not spatiotopic coordinates across several natural image transformations. Journal of vision 9, 10.1-1017.

[36] Rao, R. P. and Ballard, D. H. (1999). Predictive coding in the visual cortex: a functional interpretation of some extra-classical receptive-field effects. Nature neuroscience 2, 79–87.

[37] Issa, E. B.; Cadieu, C. F. and DiCarlo, J. J. (2018). Neural dynamics at successive stages of the ventral visual stream are consistent with hierarchical error signals. eLife 7, e42870.

[38] Keller, G. B. and Mrsic-Flogel, T. D. (2018). Predictive Processing: A Canonical Cortical Computation. Neuron 100, 424–435.

[39] Vinken, K.; Boix, X. and Kreiman, G. (2020). Incorporating intrinsic suppression in deep neural networks captures dynamics of adaptation in neurophysiology and perception Science Advances 6, eabd4205.

[40] Chaudhuri, R.; Knoblauch, K.; Gariel, M.-A.; Kennedy, H. and Wang, X.-J. (2015). A Large-Scale Circuit Mechanism for Hierarchical Dynamical Processing in the Primate Cortex. Neuron 88, 419–431.

[41] Himberger, K. D.; Chien, H.-Y. and Honey, C. J. (2018). Principles of Temporal Processing Across the Cortical Hierarchy. Neuroscience 389, 161–174.

[42] Murray, J. D.; Bernacchia, A.; Freedman, D. J.; Romo, R.; Wallis, J. D.; Cai, X.; Padoa-Schioppa, C.; Pasternak, T.; Seo, H.; Lee, D. and Wang, X.-J. (2014). A hierarchy of intrinsic timescales across primate cortex. Nature neuroscience 17, 1661–1663.

[43] Runyan, C. A.; Piasini, E.; Panzeri, S. and Harvey, C. D. (2017). Distinct timescales of population coding across cortex. Nature 548, 92–96.

[44] Niell, C. M. and Stryker, M. P. (2008). Highly selective receptive fields in mouse visual cortex. The Journal of neuroscience: the official journal of the Society for Neuroscience 28, 7520–7536.

[45] Cohen, M. R. and Maunsell, J. H. R. (2009). Attention improves performance primarily by reducing interneuronal correlations Nature Neuroscience 12, 1594–1600.

[46] Ayaz, A.; Saleem, A. B.; Schölvinck, M. L. and Carandini, M. (2013). Locomotion Controls Spatial Integration in Mouse Visual Cortex Current Biology 23, 890–894.

[47] Froudarakis, E.; Berens, P.; Ecker, A. S.; Cotton, R. J.; Sinz, F. H.; Yatsenko, D.; Saggau, P.; Bethge, M. and Tolias, A. S. (2014). Population code in mouse V1 facilitates readout of natural scenes through increased sparseness. Nature neuroscience 17, 851–857.

[48] Burgess, C. R.; Ramesh, R. N.; Sugden, A. U.; Levandowski, K. M.; Minnig, M. A.; Fenselau, H.; Lowell, B. B. and Andermann, M. L. (2016). Hunger-Dependent Enhancement of Food Cue Responses in Mouse Postrhinal Cortex and Lateral Amygdala. Neuron 91, 1154–1169.

[49] Khan, A. G. and Hofer, S. B. (2018). Contextual signals in visual cortex Current Opinion in Neurobiology 52, 131–138.

[50] Lehky, S. R. and Sereno, A. B. (2007). Comparison of Shape Encoding in Primate Dorsal and Ventral Visual Pathways Journal of Neurophysiology 97, 307–319.

[51] Glickfeld, L. L. and Olsen, S. R. (2017). Higher-Order Areas of the Mouse Visual Cortex Annual Review of Vision Science 3, 251–273.

[52] Glickfeld, L. L.; Reid, R. C. and Andermann, M. L. (2014). A mouse model of higher visual cortical function Current Opinion in Neurobiology 24, 28–33.

[53] Tafazoli, S.; Safaai, H.; De Franceschi, G.; Rosselli, F. B.; Vanzella, W.; Riggi, M.; Buffolo, F.; Panzeri, S. and Zoccolan, D. (2017). Emergence of transformation-tolerant representations of visual objects in rat lateral extrastriate cortex. eLife 6, e22794.

[54] Vermaercke, B.; Gerich, F. J.; Ytebrouck, E.; Arckens, L.; Op de Beeck, H. P. and Van den Bergh, G. (2014). Functional specialization in rat occipital and temporal visual cortex Journal of neurophysiology 112, 1963–1983.

[55] Matteucci, G.; Bellacosa Marotti, R.; Riggi, M.; Rosselli, F. B. and Zoccolan, D. (2019). Nonlinear Processing of Shape Information in Rat Lateral Extrastriate Cortex. The Journal of neuroscience: the official journal of the Society for Neuroscience 39, 1649–1670.

[56] Froudarakis, E.; Cohen, U.; Diamantaki, M.; Walker, E. Y.; Reimer, J.; Berens, P.; Sompolinsky, H. and Tolias, A. S. (2020). Object manifold geometry across the mouse cortical visual hierarchy. biorxiv preprint. doi:10.1101/2020.08.20.258798

[57] Matteucci, G. and Zoccolan, D. (2020). Unsupervised experience with temporal continuity of the visual environment is causally involved in the development of V1 complex cells. Science Advances 6, eaba3742.

[58] Tafazoli, S.; Filippo, A. D. and Zoccolan, D. (2012). Transformation-Tolerant Object Recognition in Rats Revealed by Visual Priming Journal of Neuroscience 32, 21–34.

[59] Vinken, K.; Van den Bergh, G.; Vermaercke, B. and Op de Beeck, H. P. (2016). Neural Representations of Natural and Scrambled Movies Progressively Change from Rat Striate to Temporal Cortex. Cerebral cortex (New York, N.Y.: 1991) 26, 3310–3322.

[60] Siegle, J. H.; Jia, X.; Durand, S.; Gale, S.; Bennett, C.; Graddis, N.; Heller, G.; Ramirez, T. K.; Choi, H.; Luviano, J. A.; Groblewski, P. A.; Ahmed, R.; Arkhipov, A.; Bernard, A.; Billeh, Y. N.; Brown, D.; Buice, M. A.; Cain, N.; Caldejon, S.; Casal, L.; Cho, A.; Chvilicek, M.; Cox, T. C.; Dai, K.; Denman, D. J.; de Vries, S. E. J.; Dietzman, R.; Esposito, L.; Farrell, C.; Feng, D.; Galbraith, J.; Garrett, M.; Gelfand, E. C.; Hancock, N.; Harris, J. A.; Howard, R.; Hu, B.; Hytnen, R.; Iyer, R.; Jessett, E.; Johnson, K.; Kato, I.; Kiggins, J.; Lambert, S.; Lecoq, J.; Ledochowitsch, P.; Lee, J. H.; Leon, A.; Li, Y.; Liang, E.; Long, F.; Mace, K.; Melchior, J.; Millman, D.; Mollenkopf, T.; Nayan, C.; Ng, L.; Ngo, K.; Nguyen, T.; Nicovich, P. R.; North, K.; Ocker, G. K.; Ollerenshaw, D.; Oliver, M.; Pachitariu, M.; Perkins, J.; Reding, M.; Reid, D.; Robertson, M.; Ronellenfitch, K.; Seid, S.; Slaughterbeck, C.; Stoecklin, M.; Sullivan, D.; Sutton, B.; Swapp, J.; Thompson, C.; Turner, K.; Wakeman, W.; Whitesell, J. D.; Williams, D.; Williford, A.; Young, R.; Zeng, H.; Naylor, S.; Phillips, J. W.; Reid, R. C.; Mihalas, S.; Olsen, S. R. and Koch, C. (2019). A survey of spiking activity reveals a functional hierarchy of mouse corticothalamic visual areas. biorxiv preprint. doi:10.1101/805010

[61] Espinoza, S. G. and Thomas, H. C. (1983). Retinotopic organization of striate and extrastriate visual cortex in the hooded rat Brain Research 272, 137–144.

[62] Fiser, J.; Chiu, C. and Weliky, M. (2004). Small modulation of ongoing cortical dynamics by sensory input during natural vision. Nature 431, 573–578.

[63] Averbeck, B. B.; Latham, P. E. and Pouget, A. (2006). Neural correlations, population coding and computation. Nature reviews. Neuroscience 7, 358–366.

[64] Valente, M.; Pica, G.; Runyan, C. A.; Morcos, A. S.; Harvey, C. D. and Panzeri, S. (2020). Correlations enhance the behavioral readout of neural population activity in association cortex. doi:10.1101/2020.04.03.024133

[65] Vascon, S.; Parin, Y.; Annavini, E.; D’Andola, M.; Zoccolan, D. and Pelillo, M. (2018). Characterization of Visual Object Representations in Rat Primary Visual Cortex. The European Conference on Computer Vision (ECCV) Workshops,.

[66] Harris, J. A.; Mihalas, S.; Hirokawa, K. E.; Whitesell, J. D.; Choi, H.; Bernard, A.; Bohn, P.; Caldejon, S.; Casal, L.; Cho, A.; Feiner, A.; Feng, D.; Gaudreault, N.; Gerfen, C. R.; Graddis, N.; Groblewski, P. A.; Henry, A. M.; Ho, A.; Howard, R.; Knox, J. E.; Kuan, L.; Kuang, X.; Lecoq, J.; Lesnar, P.; Li, Y.; Luviano, J.; McConoughey, S.; Mortrud, M. T.; Naeemi, M.; Ng, L.; Oh, S. W.; Ouellette, B.; Shen, E.; Sorensen, S. A.; Wakeman, W.; Wang, Q.; Wang, Y.; Williford, A.; Phillips, J. W.; Jones, A. R.; Koch, C. and Zeng, H. (2019). Hierarchical organization of cortical and thalamic connectivity Nature 575, 195–202.

[67] Wang, Q.; Gao, E. and Burkhalter, A. (2011). Gateways of Ventral and Dorsal Streams in Mouse Visual Cortex Journal of Neuroscience 31, 1905–1918.

[68] Wang, Q.; Sporns, O. and Burkhalter, A. (2012). Network Analysis of Corticocortical Connections Reveals Ventral and Dorsal Processing Streams in Mouse Visual Cortex Journal of Neuroscience 32, 4386–4399.

[69] Gilbert, C. D. and Li, W. (2013). Top-down influences on visual processing. Nature reviews. Neuroscience 14, 350–363.

[70] Homann, J.; Koay, S. A.; Glidden, A. M.; Tank, D. W. and Berry, M. J. (2017). Predictive Coding of Novel versus Familiar Stimuli in the Primary Visual Cortex. doi:10.1101/197608

[71] Liao, Q. and Poggio, T. (2016). Bridging the Gaps Between Residual Learning, Recurrent Neural Networks and Visual Cortex. arxiv preprint. doi:

[72] Kubilius, J.; Schrimpf, M.; Nayebi, A.; Bear, D.; Yamins, D. L. K. and DiCarlo, J. J. (2018). CORnet: Modeling the Neural Mechanisms of Core Object Recognition. biorxiv preprint. doi:10.1101/408385

[73] Tang, H.; Schrimpf, M.; Lotter, W.; Moerman, C.; Paredes, A.; Ortega Caro, J.; Hardesty, W.; Cox, D. and Kreiman, G. (2018). Recurrent computations for visual pattern completion. Proceedings of the National Academy of Sciences of the United States of America 115, 8835–8840.

[74] Kar, K.; Kubilius, J.; Schmidt, K.; Issa, E. B. and DiCarlo, J. J. (2019). Evidence that recurrent circuits are critical to the ventral stream’s execution of core object recognition behavior. Nature neuroscience 22, 974–983.

[75] Kietzmann, T. C.; Spoerer, C. J.; Sörensen, L. K. A.; Cichy, R. M.; Hauk, O. and Kriegeskorte, N. (2019). Recurrence is required to capture the representational dynamics of the human visual system Proceedings of the National Academy of Sciences 116, 21854–21863.

[76] Laughlin, S. (1981). A simple coding procedure enhances a neuron’s information capacity. Zeitschrift fur Naturforschung. Section C, Biosciences 36, 910–912.

[77] Tkacik, G.; Prentice, J. S.; Victor, J. D. and Balasubramanian, V. (2010). Local statistics in natural scenes predict the saliency of synthetic textures. Proceedings of the National Academy of Sciences of the United States of America 107, 18149–18154.

[78] Olshausen, B. A. and Field, D. J. (1996). Emergence of simple-cell receptive field properties by learning a sparse code for natural images. Nature 381, 607–609.

[79] Li, Y.; Fitzpatrick, D. and White, L. E. (2006). The development of direction selectivity in ferret visual cortex requires early visual experience. Nature neuroscience 9, 676–681.

[80] Li, Y.; Van Hooser, S. D.; Mazurek, M.; White, L. E. and Fitzpatrick, D. (2008). Experience with moving visual stimuli drives the early development of cortical direction selectivity. Nature 456, 952–956.

[81] Hunt, J. J.; Dayan, P. and Goodhill, G. J. (2013). Sparse coding can predict primary visual cortex receptive field changes induced by abnormal visual input. PLoS computational biology 9, e1003005.

[82] Hermundstad, A. M.; Briguglio, J. J.; Conte, M. M.; Victor, J. D.; Balasubramanian, V. and Tkačik, G. (2014). Variance predicts salience in central sensory processing. eLife 3, e03722.

[83] Kuang, X.; Poletti, M.; Victor, J. D. and Rucci, M. (2012). Temporal encoding of spatial information during active visual fixation. Current biology: CB 22, 510–514.

[84] Kleiner, M.; Brainard, D.; Pelli, D.; Ingling, A.; Murray, R. and Broussard, C. (2007). What’s new in psychtoolbox-3? Perception 36, 1–16.

[85] Marshel, J. H.; Garrett, M. E.; Nauhaus, I. and Callaway, E. M. (2011). Functional specialization of seven mouse visual cortical areas. Neuron 72, 1040–1054.

[86] Rossant, C.; Kadir, S. N.; Goodman, D. F. M.; Schulman, J.; Hunter, M. L. D.; Saleem, A. B.; Grosmark, A.; Belluscio, M.; Denfield, G. H.; Ecker, A. S.; Tolias, A. S.; Solomon, S.; Buzsaki, G.; Carandini, M. and Harris, K. D. (2016). Spike sorting for large, dense electrode arrays. Nature neuroscience 19, 634–641.

[87] Rikhye, R. V. and Sur, M. (2015). Spatial Correlations in Natural Scenes Modulate Response Reliability in Mouse Visual Cortex. The Journal of neuroscience: the official journal of the Society for Neuroscience 35, 14661–14680.

[88] Katsnelson, J. and Kotz, S. (1957). On the upper limits of some measures of variability Archiv für Meteorologie, Geophysik und Bioklimatologie Serie B 8, 103–107.

[89] Wales, D. J. and Scheraga, H. A. (1999). Global Optimization of Clusters, Crystals, and Biomolecules Science 285, 1368–1372.

[90] Byrd, R. H.; Lu, P.; Nocedal, J. and Zhu, C. (1995). A Limited Memory Algorithm for Bound Constrained Optimization SIAM Journal on Scientific Computing 16, 1190–1208.

[91] Bates, D. M. and Watts, D. G. (1988). Nonlinear regression analysis and its applications. Wiley New York.

[92] Fan, R.-E.; Chang, K.-W.; Hsieh, C.-J.; Wang, X.-R. and Lin, C.-J. (2008). LIBLINEAR: A Library for Large Linear Classification J. Mach. Learn. Res. 9, 1871–1874.

[93] Pedregosa, F.; Varoquaux, G.; Gramfort, A.; Michel, V.; Thirion, B.; Grisel, O.; Blondel, M.; Prettenhofer, P.; Weiss, R.; Dubourg, V.; Vanderplas, J.; Passos, A.; Cournapeau, D.; Brucher, M.; Perrot, M. and Duchesnay, E. (2011). Scikit-learn: Machine Learning in Python Journal of Machine Learning Research 12, 2825–2830.

[94] Wilcox, R. R. (2011). Introduction to robust estimation and hypothesis testing. Academic press.

[95] Efron, B. and Tibshirani, R. J. (1994). An introduction to the bootstrap. CRC press.

[96] R Core Team (2020). R: A Language and Environment for Statistical Computing,.

[97] Vinken, K. (2020). Data from “Neural representations of natural and scrambled movies progressively change from rat striate to temporal cortex” doi:10.17605/OSF.IO/M2E6D

