## Supplementary Information for "Intrinsic dynamics enhance temporal stability of stimulus representation along rodent visual cortical hierarchies"

### Supplementary Figures

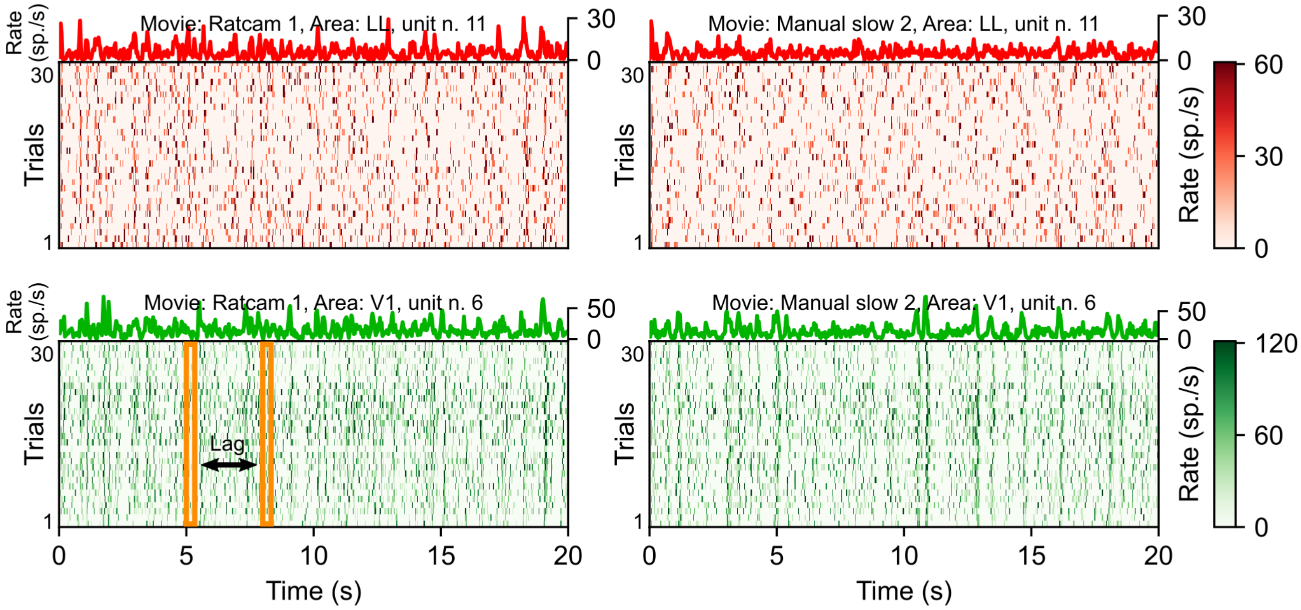


**Supplementary Fig. 1 Schematic illustrating the computation of intrinsic correlations.** Each panel shows the binned firing rate of an example neuron in response to a given movie across 30 repeated presentations (or trials) of the stimulus. Top: representative neuron in LL. Bottom: representative neuron in V1. Left: ratcam 1 movie. Right: manual slow 2 movie. By analogy to the display in Fig. 3c, the top part of each item gives the trial-averaged firing rate (PSTH) for the chosen neuron. For concreteness, the 11$^{th}$ row of the top-left matrix in Fig. 3c contains the same data as the red line shown here in the top-left panel. The yellow boxes indicate the vectors over which intrinsic correlations are computed. Note that here the vectors contain the (binned) activity of a single neuron over multiple trials, while in Fig. 3c (where signal correlations were discussed) the vectors contained the trial-averaged activity of multiple cells within an area.

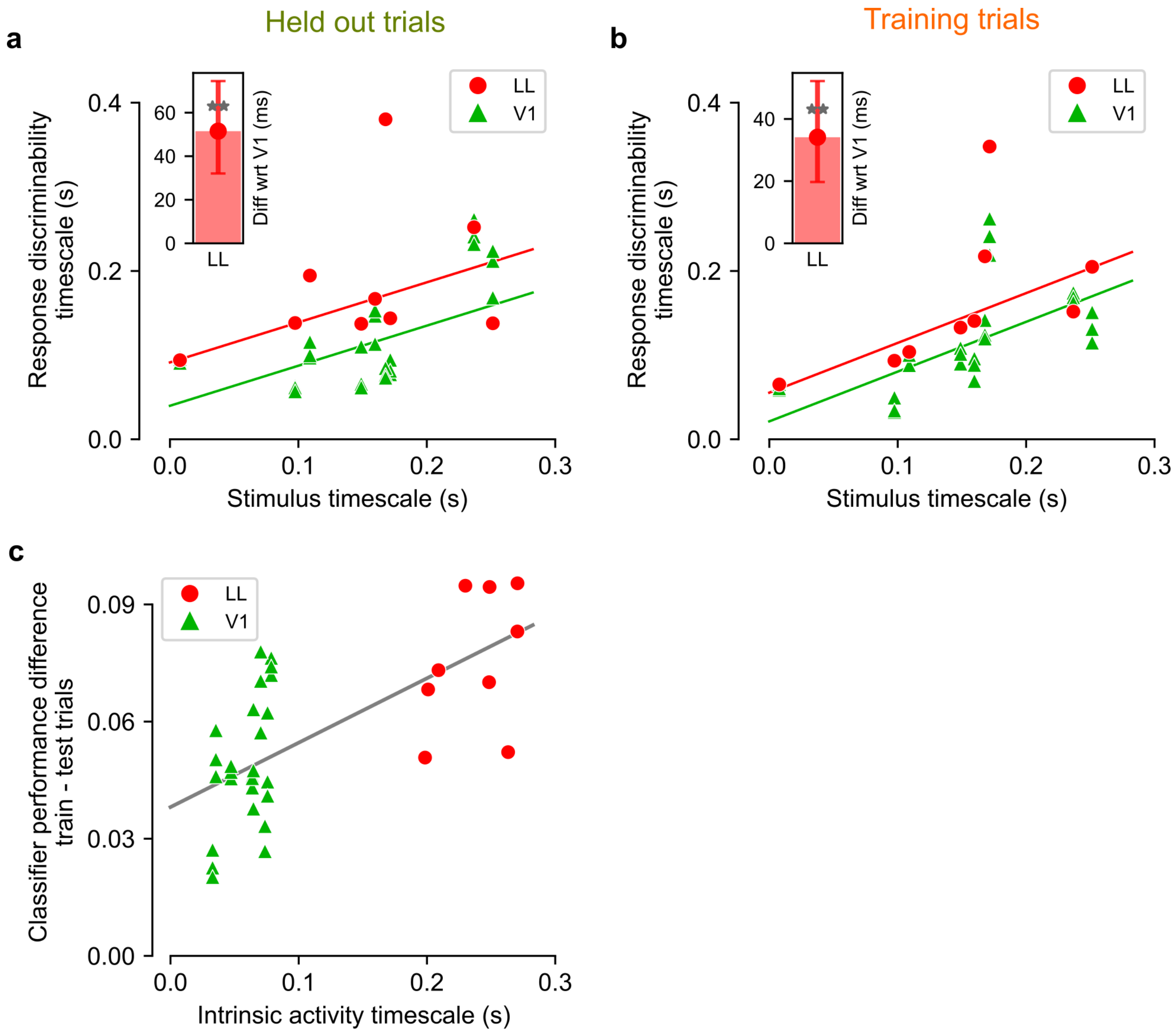


**Supplementary Fig. 2 Decoding analysis for larger pseudopopulations.** The pseudopopulation size we selected for our analyses (K = 20) was dictated by the need for having at least one pseudopopulation for each area we recorded from. By restricting our analyses to V1 and LL only (i.e., the areas that were more densely sampled in our recordings), we were able to confirm that our results hold for larger populations of 50 units. **a** The timescale of response discriminability, measured as the time constant of the classifier performance on held-out trials (i.e., green boxes in Fig. 5a), is plotted as a function of the timescale of the movies. Each dot corresponds to a distinct pseudopopulation. The solid lines are linear regressions (Theil-Sen robust estimator) with common slope and different intercept across the two areas. The inset shows the difference of the intercept between area LL and V1 (**, p=3e-3, one-tailed bootstrap test). Slope: 0.48, 68%CI [0.29-0.66] (percentile bootstrap, see Methods). V1 intercept: 40[16-68] ms. Intercept difference LL-V1: 52[32-75] ms. This analysis is equivalent to that shown in Fig. 5c, but using pseudopopulations of 50 units instead of 20. **b** Same as **a**, but for the timescale of response discriminability measured on training trials (i.e., orange boxes in Fig. 5a). Slope: 0.59[0.46-0.72]. V1 intercept: 21[-1-44] ms. Intercept difference LL-V1: 34[20-52] ms. As shown in the inset, such difference was significantly larger than zero (**, p=2e-3, one-tailed bootstrap test). This analysis is equivalent to that shown in Supplementary Fig. 3b, but using pseudopopulations of 50 units instead of 20. **c** The amount of classifier performance due to intrinsic correlations is plotted as a function of the timescale of such correlations (i.e., the intrinsic timescale of neuronal activity shown in Fig. 4d). This analysis is equivalent to that shown in Fig. 5d, but using pseudopopulations of 50 units instead of 20. The gray line represents a linear fir to the data. Slope of the fit: 0.17±0.03, nonzero p=2e-5, two-tailed t-test, t=5.0, df=34; intercept: 0.038±0.004, p=3e-10, two-tailed t-test, t=8.7, df=34.

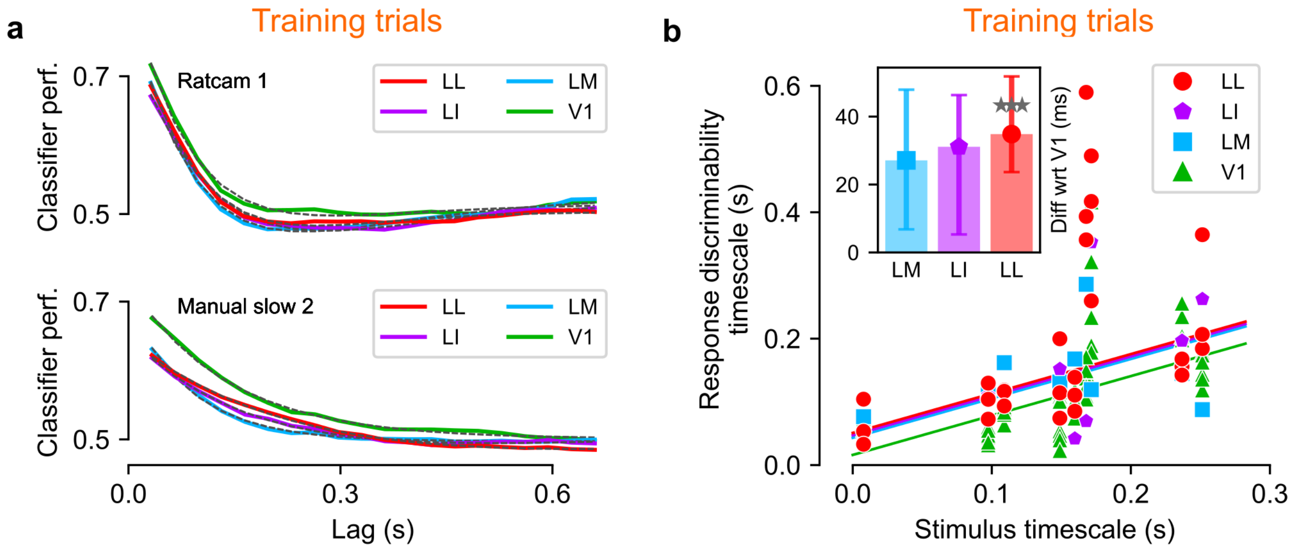


**Supplementary Fig. 3 Decoding analysis on training trial set. a** Classifier performance on the training trial set, for two example movie stimuli and one example pseudopopulation per area (colored curves). The performance is plotted as a function of the lag between training bin (i.e., gray boxes in Fig. 5a) and test bin (i.e., orange boxes in Fig. 5a), and is fitted with either an exponential decay or a damped oscillation function (fits are shown as black dashed lines). These curves are equivalent to those shown in Fig. 5b, but they were obtained by computing classification performances on the same trials used to train the classifier (i.e., orange boxes in Fig. 5a), rather than on held-out trials (i.e., green boxes in Fig. 5a). **b** The timescale of response discriminability, measured as the time constant of the exponential decay of the classifier performance on training trials, is plotted as a function of the timescale of the movies. Each dot corresponds to a distinct pseudopopulation. The solid lines are linear regressions with common slope and different intercept across the four areas. The inset shows the difference of the intercept for areas LM, LI and LL vs area V1. Common slope: 0.62, 68%CI [0.55-0.70], percentile bootstrap. V1 intercept: 16[2-29] ms. The intercept of the fit was significantly different for LL vs. V1 ($p$=0.1, 0.12, 3e-4 respectively for LM, LI, LL, one-tailed bootstrap test). Intercept differences were: LM 27[7-48] ms; LI 31[5-46] ms; LL 35[24-52] ms.

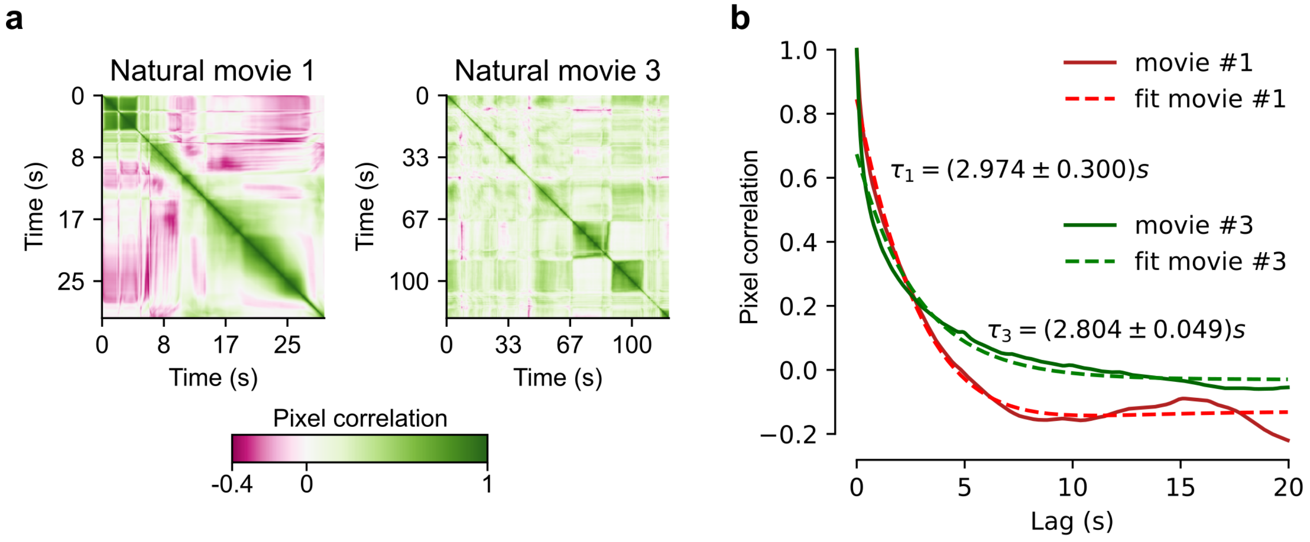


**Supplementary Fig. 4 Pixel correlations for the movies in the Allen stimulus set. a** Pixel correlation matrix, showing the correlation coefficient between each pair of frames, for both movies in the stimulus set. Note the strong block structure in Natural Movie 1. **b** Pixel correlation functions obtained by averaging over the diagonals of the matrices in **a**. Solid lines are the empirical correlation functions, dashed lines of corresponding color are exponential fits. The extracted timescales are reported. Natural Movie 1 was discarded due to the presence of the heavy irregular tail visible in **b**, due to the block structure highlighted in **a**.

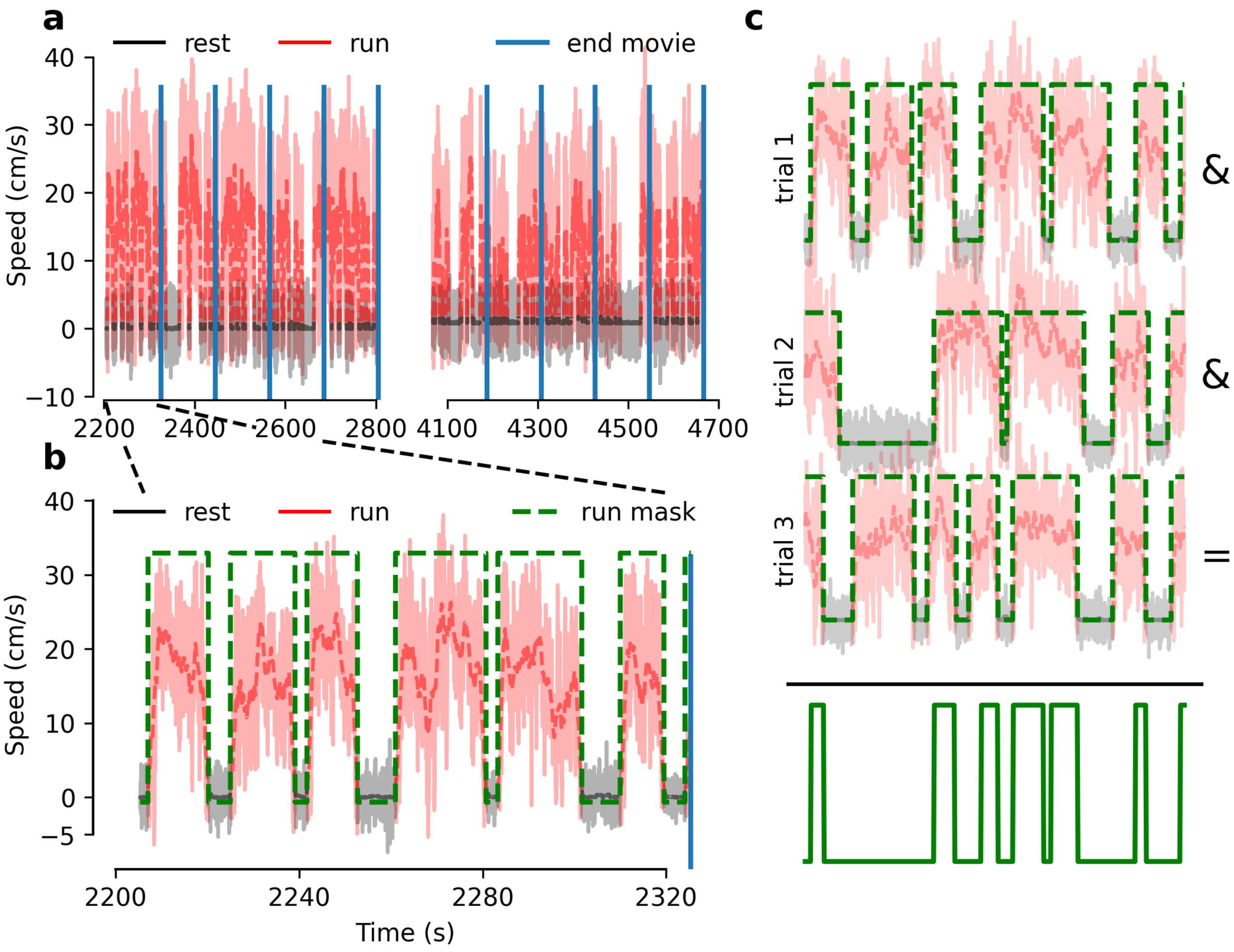


**Supplementary Fig. 5 Identification of running and resting state. a** Example of spinning wheel velocity during the presentation of Natural Movie 3 from the first session in the Allen dataset (2 blocks of 5 trials each). The running condition is identified as v > 1cm/s and is colored in red, resting is in gray. Solid lines are raw values, dashed lines are the corresponding smoothed averages (window size 1.7s). Solid vertical blue lines marks the end of a given trial. **b** Zoomed plot of first trial. Overlaid green dashed line is the mask associated to the running state for this given trial and session. Vertical line marks the end of the trial. **c** Example of the computation of the intersection between the first three trials of the example session. The resulting mask, bottom solid green line, selects the times of shared running state between the trials under exam.

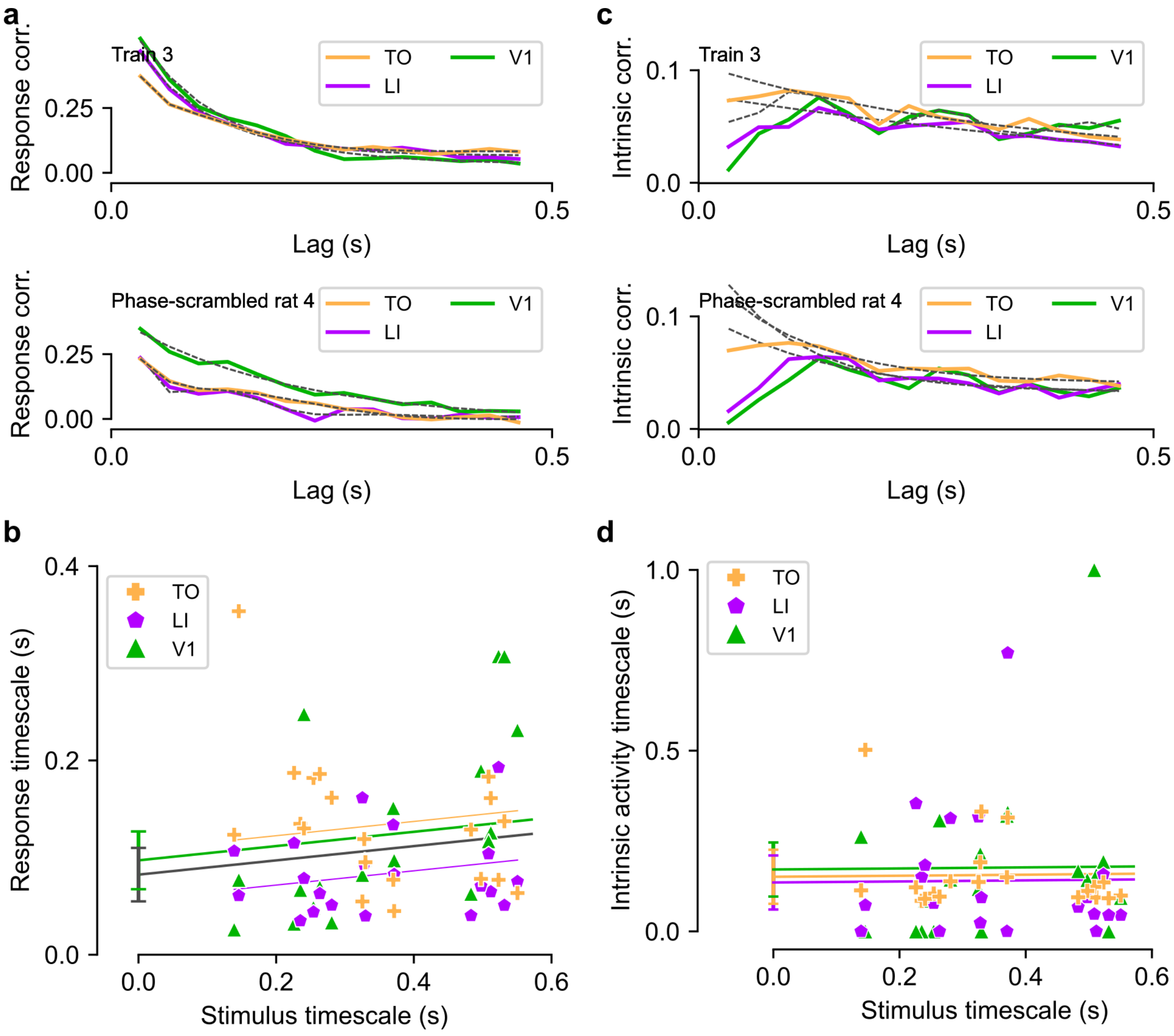


**Supplementary Fig. 6 Correlation analysis for awake rat data.** As Fig. 4, for the data recorded in awake rat. **a** Correlation between trial-averaged population vectors as a function of the temporal lag separating them (examples for two representative movies). Colored lines: empirical data. Dashed lines: fits. **b** Population response timescales as a function of the timescales of the corresponding movie stimuli (colored markers). Each colored line is a linear fit prediction of the relationship between such timescales for a given area (same color code as in the key). The gray line is the linear fit prediction obtained by pooling together the data of the two extrastriate areas (i.e., LI and LL). The error bars show the standard errors of the intercept of the linear fits for V1 and the pooled extrastriate areas. **c** and **d** Same as **a** and **b**, but for the intrinsic timescales of neuronal activity.

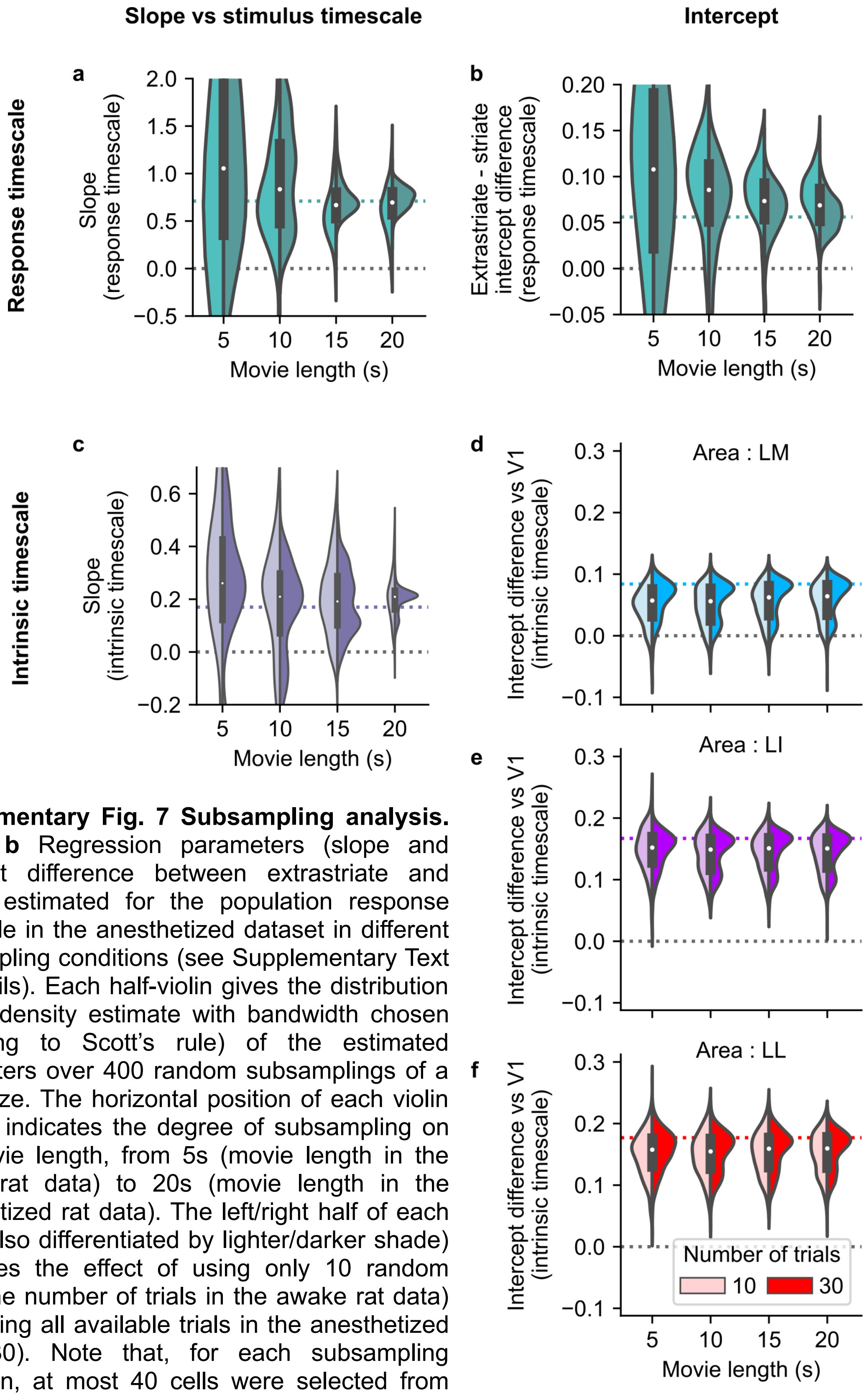


**Supplementary Fig. 7 Subsampling analysis.** **a** and **b** Regression parameters (slope and intercept difference between extrastriate and striate) estimated for the population response timescale in the anesthetized dataset in different subsampling conditions (see Supplementary Text for details). Each half-violin gives the distribution (kernel density estimate with bandwidth chosen according to Scott's rule) of the estimated parameters over 400 random subsamplings of a given size. The horizontal position of each violin (x axis) indicates the degree of subsampling on the movie length, from 5s (movie length in the awake rat data) to 20s (movie length in the anesthetized rat data). The left/right half of each violin (also differentiated by lighter/darker shade) compares the effect of using only 10 random trials (the number of trials in the awake rat data) vs keeping all available trials in the anesthetized data (30). Note that, for each subsampling repetition, at most 40 cells were selected from each cortical area to facilitate the comparison with the awake rat data. Colored dashed lines: estimates on full dataset reported in the paper. **c-f** as **a-b**, for the intrinsic timescales.

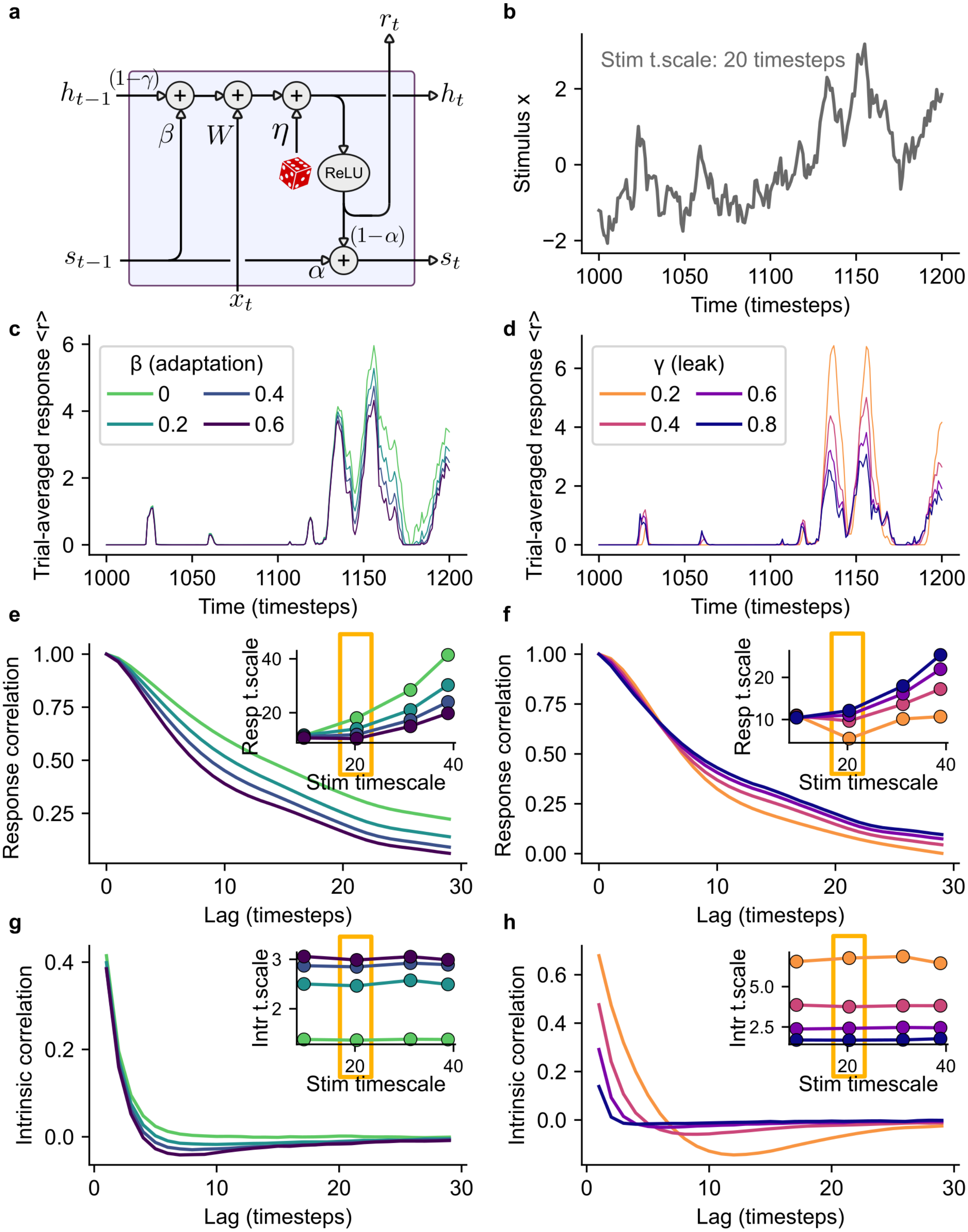

a
b
c
d
e
f
g
h
ReLU
Stim t.scale: 20 timesteps
Stimulus x
Time (timesteps)
Trial-averaged response <r>
β (adaptation)
γ (leak)
Response correlation
Resp t.scale
Stim timescale
Lag (timesteps)
Intrinsic correlation
Intr t.scale

**Supplementary Fig. 8 Conceptual model illustrating the relationship between adaptive and intrinsic processing and activity timescales. a** Schematic representation of model dynamics (see equations in Supplementary Text). **b** Example time course for the total synaptic input x. The stimulus and the systems were simulated for 3000 timesteps; here about 200 timesteps are shown for illustrative purposes. The stimulus shown was generated as a Gaussian process with absolute exponential kernel with a timescale 20 timesteps. Other stimulus instances were simulated, with nominal timescales of 10, 30, and 40 timesteps. **c** Example PSTH (average response over 1000 trials) of the simulated system in response to the stimulus instance with a nominal timescale of 20 timestep (shown in panel **b**) for different values of the adaptation strength parameter, keeping all other parameters fixed (in particular, $\gamma$=0.5). **e** Response correlation function (autocorrelation of the PSTH) for the simulated conditions in panel **c**. Inset: estimated response timescale as a function of stimulus timescale and adaptation strength. The yellow box highlights the condition shown in the main panel (stimulus with 20-timesteps timescale). **g** As **e**, for intrinsic correlations (correlations of the simulated activity across trials). **d, f, h** As **c, g, f**, when varying the leak rate parameter of the simulated system and keeping all other parameters fixed (in particular, $\beta$=0.6).

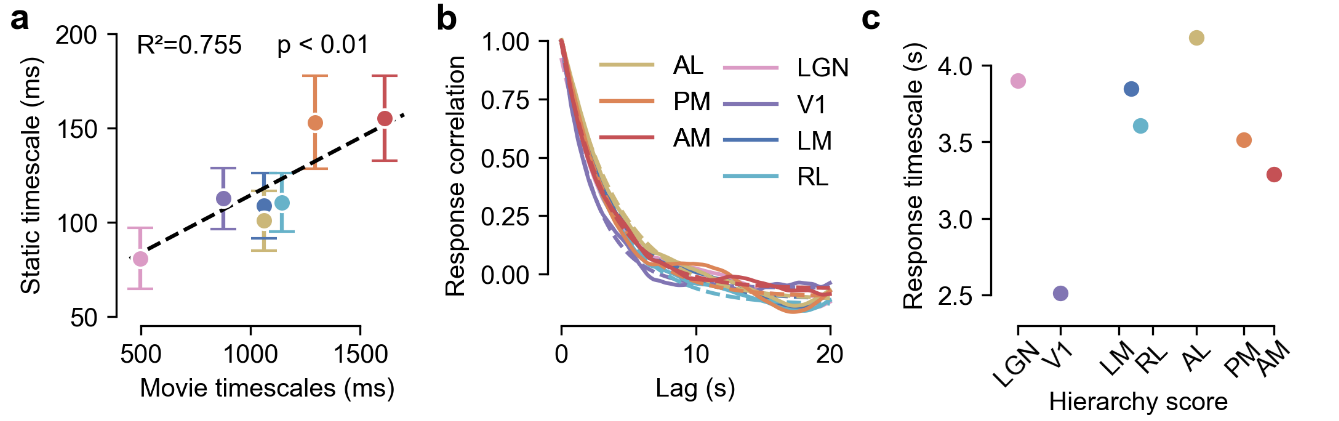


**Supplementary Fig. 9 Linear Model for Allen data. a** y-axis: timescale of the spike autocorrelation during response to visual flash (see Discussion). x-axis: response timescale during movie presentation. **b** Response correlation of the linear model when simulated with the movie stimulus. Solid lines: empirical correlations. Dashed lines: exponential fits. **c** Response timescales computed from the linear model response, displayed versus the anatomical hierarchy score of each area.

### Supplementary text

#### Subsampling analysis of anesthetized data

In order to assess the extent to which the analysis of awake, body-constrained rat data (Supplementary Fig. 6) is expected to be informative regarding the hypotheses studied elsewhere in the paper, we performed a subsampling analysis of our core dataset recorded in anesthetized rat. The awake dataset consisted in recordings performed in three cortical areas, with about 40 single-units per area (V1: 44, LI: 40, TO: 38), in response to 20 movies, each of which was 5 seconds long and repeated 10 times. By contrast, the anesthetized dataset contained data from four cortical areas, with a less-uniform distribution of units per area (V1: 168, LM: 20, LI: 36, LL: 70), recorded in response to 9 movies, each of which was 20 seconds long and repeated 30 times. We therefore sought to understand how the results from the anesthetized data would look if we limited the experimental data to a comparable amount to that recorded in awake rats, especially with respect to the number of trials and movie duration. To do this, we repeated the analyses reported in Fig. 4 a number of times, each time sampling a random subset of the available data. More in detail, for a given, desired, number of trials and movie duration, we proceeded as follows:

- For each movie, we selected a contiguous segment of the movie of the desired duration, taken at random.
- We computed the stimulus timescale based on that random segment.
- We discarded all neural data recorded outside of that movie segment.
- We selected a random subset of trials of the desired size, and discarded all other trials from the neural recordings.
- For each area, we selected a random neural subpopulation of at most 40 cells (or all available cells when these are less than 40), and discarded all others.
- On this subsampled neural dataset, we computed the response and intrinsic time scale for each area and each movie (this involved computing the empirical correlation functions, fitting them with our model selection procedure, and extracting the timescale parameter).
- We regressed the response timescale vs stimulus timescale and "macro area" (striate | extra-striate), as in Fig. 4b.
- We regressed the intrinsic timescale vs stimulus timescale and area, as in Fig. 4d.
- For both regression analyses, we extract a slope parameter and a parameter associated with "intercept difference with respect to V1", which for the response timescale is simply one number (extrastriate-striate intercept difference) and for the intrinsic timescale is one number per extrastriate area (LM-V1, LI-V1, LL-V1).
- We repeated all points above 400 times, and computed the resulting distributions over the slope and intercept parameters.

Finally, we visualized the distribution of the regression parameters (slope, intercept) for both types of timescales (response, intrinsic) as a function of the number of trials and movie duration used (Supplementary Fig. 7). As mentioned in the main text, in the subsampling condition that most closely approximates the awake dataset (movie duration 5 seconds, 10 trials per movie) the distributions of some of these parameters straddle the null result (zero) with a very large portion of their mass, suggesting that the analysis of the awake data is not expected to provide statistically significant results, just by lack of statistical power, if the underlying effects had the same strength they have in the anesthetized data. More specifically, we do not expect to see any significant effect for either the slope or the intercept term of the response timescales, nor for the slope of the intrinsic timescales. On the other hand, the intercept term of the intrinsic timescales, capturing the difference of the intrinsic timescale in each of the extrastriate areas versus V1, seems fairly robust to subsampling and would thus be expected to yield significant results in the awake data if these were roughly homogeneous to the anesthetized data, and in particular if brain state did not affect correlations. Therefore, the fact that intrinsic timescales do not show any difference across areas in the awake dataset (where the rats were forced in a resting condition) supports the conclusion drawn from the Allen dataset, that this difference across areas is highly dependent on the state of the animal, and specifically it is weaker during quiet wakefulness compared to active behavior.

### A conceptual model illustrating the relationship between adaptive and intrinsic mechanisms and activity timescales

#### Model definition

In this section, we give some proof-of-principle examples of how intrinsic and adaptive dynamical processes can affect neural activity timescales, at both the single-trial and trial-averaged level, expanding on the ideas illustrated in the cartoons of Fig. 1. To do so, we make use of a simple model that we obtained by generalizing the model presented in [1] to include integrator-like dynamics and internal noise. Briefly, we consider an idealized setting where a time-varying sensory stimulus of a certain duration is presented to an animal for a certain number of trials. At each trial, the firing rate $\mathrm{r}_t$ of a neuron is recorded (as a function of time $\mathrm{t}$). The temporal evolution of the neural response is modeled according to the following set of equations:

$$s_t = \alpha s_{t-1} + (1-\alpha) r_t \qquad \text{(Equation .1)}$$

$$h_t = b + W x_t - \beta s_t + (1-\gamma) h_{t-1} + \eta \qquad \text{(Equation .2)}$$

$$r_t = \sigma(h_t) \qquad \text{(Equation .3)}$$

where $\mathrm{x}_t$ represents the net synaptic input to the neuron at time $\mathrm{t}$, $\mathrm{s}_t$ is an auxiliary "state" variable used to implement intrinsic suppression (see [1] for details), $\eta$ is a noise term sampled independently at each time step (we will assume $\eta \sim \mathcal{N}(0, \rho)$ for some fixed standard deviation $\rho$), $\sigma(\cdot)$ is the ReLU nonlinearity, and $\alpha, \beta, \gamma, b, W$ are parameters. The model can be represented graphically as in Supplementary Fig. 8a, and the parameters can be given the following intuitive meanings: $\mathrm{b}$ is a bias term, $\mathrm{W}$ is a synaptic weigth, $\alpha$ controls the integration time constant of the state variable, $\beta$ is the adaptation strength (i.e., the strength of the intrinsic suppression mechanism), and $\gamma$ is related to the leak rate of the system seen, when $\beta = 0$, as a leaky integrator (or its cutoff frequency, if seen as a low-pass filter). This model reduces to that in [1] if we set $\rho = 0$ and $\gamma = 1$; in other words, this model extends that presented in [1] by adding an internal noise term that is sampled independently at each timestep of each trial and by allowing the system to integrate its inputs over time. On the other hand, we note that what we present here is simpler than the model discussed in [1] because only one neuron is modeled (rather than a network), and therefore all quantities appearing in the equations above are scalars. For simplicity, in the following we will fix $\mathrm{b}=0$, $\mathrm{W}=1$, $\rho = 0.3$ and $\alpha = 0.96$ (this last value being the one used throughout [1]) unless specified otherwise.

#### Stimulus details and simulation set-up

In our simulations, we generated the input $\mathrm{x}_t$ according to a Gaussian process with absolute exponential kernel with a certain timescale $\tau_s$ (see example in Supplementary Fig. 8b). In other words, a stimulus of length T was generated by taking a sample from a T-variate Gaussian with zero mean and covariance matrix $\Sigma_{i,j} = \exp[-\frac{|i-j|}{\tau_s}]$ (note that, for the purpose of this simulation, time is measured in units of discrete time steps, so *i-j* can be interpreted as a time difference). This allowed us to avoid building too much structure into our stimuli, while still being able to control their characteristic timescales: indeed, by construction, the empirical timescale of such a stimulus (measured with the methods used in the rest of the paper) will converge to the nominal value of $\tau_s$ when T is large.

In the simulation presented here, the duration T was fixed to 3000 timesteps. We generated four different stimuli, with nominal timescales $\tau_s$ equal to 10, 20, 30 and 40 timesteps. The empirical timescales of the stimuli (measured by fitting a decaying exponential to the autocorrelation function of each stimulus) were, respectively, 9.6, 20.4, 31.3, and 38.9 timesteps. For each of these four stimuli, we simulated the response of the system over 1000 trials. For each trial, the stimulus was the same, but the internal noise of the system was sampled independently, leading to a different neural response. We used the response of the system across the 1000 trials to estimate the response correlation and intrinsic correlation of $\mathrm{r}$, and to extract the corresponding response timescales and intrinsic timescales, using the same methods we used in the rest of the paper (see Methods for details). We studied these quantities (correlations, timescales) as we varied $\beta$ and $\gamma$, the parameters of the model that control, respectively, the strength of the adaptation and the leak rate of the subthreshold integration mechanism.

#### Results

Overall, the behavior of the model changes in a complex way as a function of the parameter values, and a full qualitative investigation is out of the scope of the present paper. However, we will analyze here some particular cases that can help bolster the intuition for the elementary point we have made in Fig. 1, namely that the presence of intrinsic and adaptive mechanisms will in general affect the characteristic timescales of the system's response. The results of the simulations are shown in Supplementary Fig. 8.

In panels c, e, and g we show the effect of changing the strength of the adaptation, controlled by the parameter $\beta$. As adaptation was increased, the system tended to respond with overall shorter responses (Supplementary Fig. 8c). This was confirmed by the analysis of the correlations of the trial-averaged response, leading to the computation of the response timescale (Supplementary Fig. 8e and inset): as $\beta$ increased, the response was temporally decorrelated in the sense that the response timescale became shorter and shorter overall, and its dependence on the stimulus timescale became weaker and weaker. (For completeness, we also note that the cartoon traces in Fig. 1b are also obtained by running actual simulations of the model discussed here, with a simple boxcar-like stimulus and appropriate choice of parameters including larger $\beta$ for the dark blue trace in Fig. 1b). The presence of adaptation also affected the shape and range of intrinsic correlations, giving rise to an oscillatory component similar to that observed in experimental data (cf. Fig. 4c). For the range of parameters explored, increasing adaptation also increased the intrinsic timescales. This is consistent with the interpretation of intrinsic timescales as a characteristic time over which the system keeps memory of its past state [2; 3].

Panels d-h in Supplementary Fig. 8 show what happens if adaptation strength is kept constant but the leak rate is changed. Similarly to higher adaptation, higher leak also leads to smaller overall response (Supplementary Fig. 8d). By comparing the example PSTHs in Supplementary Fig. 8c and d, though, it is evident that the decrease in response does not happen in the same way in both cases: the shape of the response "peaks" is roughly preserved, albeit scaled, when leak is increased, while the peaks become narrower in response to an increase in adaptation. This intuition is supported quantitatively by the analysis of the activity timescales (Supplementary Fig. 8f,h). While increasing the adaptation strength decreases the response timescales and increases the intrinsic time scale, increasing the leak rate has the opposite effect: the response timescale increases (Supplementary Fig. 8f, inset) and the intrinsic timescale decreases (Supplementary Fig. 8g, inset). Intuitively, this is consistent with the idea of stronger leak leading to the response tracking the stimulus timescale more closely (thus the curves in Supplementary Fig. 8g, inset, getting closer to the identity line as leak increases), and to a shorter within-trial memory kept by the system about its past state. Finally, depending on the value of the leak rate, we can again observe the emergence of an oscillatory component to the intrinsic correlations (Supplementary Fig. 8h), similar to what we observed in real data (Fig. 4c).

In summary, the conceptual model presented here provides a concrete example of how adaptation and intrinsic mechanisms (here exemplified by the capacity of a single neuron to integrate its inputs over time, although in a network setting these could also include recurrent circuit dynamics [2; 4]) can shape the temporal structure of the neural response, by modifying the trial-averaged response (Fig. 1b, Supplementary Fig. 8c-f) and the intrinsic correlations (Supplementary Fig. 8g-h), and how state-dependent processing combined with temporally uncorrelated noise can result in temporally extended noise correlations (i.e., intrinsic correlations) [3]. In principle, this suggests that intrinsic correlations, and more generally the variability of neural responses over trials, can be a useful source of information for improving existing models of visual cortex based on convolutional neural networks [1; 5; 6], by providing independent signatures that can help constraining the space of possible mechanisms to be considered beyond simple feedforward processing. However, although our approach here is developed by generalizing some ideas from one of such state-of-the-art visual cortical models [1], a fuller investigation of visual cortical data in terms of this model goes beyond the scope of the present paper. Indeed, unlike in [1], in our discussion we have focused on an idealized neuron rather than a network arranged as a feedforward cascade. When units described by Equations 1-3 above are combined in a feedforward arrangement (or when an existing convolutional neural network is augmented so that its neurons obey those equations for certain values of the parameters), we expect the effects describe here to compound across layers and to generate rich and nontrivial behaviors extending those reported in [1].

### References

[1] Vinken, K.; Boix, X. and Kreiman, G. (2020). Incorporating intrinsic suppression in deep neural networks captures dynamics of adaptation in neurophysiology and perception *Science Advances* **6**, eabd4205.

[2] Murray, J. D.; Bernacchia, A.; Freedman, D. J.; Romo, R.; Wallis, J. D.; Cai, X.; Padoa-Schioppa, C.; Pasternak, T.; Seo, H.; Lee, D. and Wang, X.-J. (2014). A hierarchy of intrinsic timescales across primate cortex. *Nature neuroscience* **17**, 1661-1663.

[3] Runyan, C. A.; Piasini, E.; Panzeri, S. and Harvey, C. D. (2017). Distinct timescales of population coding across cortex. *Nature* **548**, 92-96.

[4] Chaudhuri, R.; Knoblauch, K.; Gariel, M.-A.; Kennedy, H. and Wang, X.-J. (2015). A Large-Scale Circuit Mechanism for Hierarchical Dynamical Processing in the Primate Cortex. *Neuron* **88**, 419-431.

[5] Yamins, D. L. K.; Hong, H.; Cadieu, C. F.; Solomon, E. A.; Seibert, D. and DiCarlo, J. J. (2014). Performance-optimized hierarchical models predict neural responses in higher visual cortex. *Proceedings of the National Academy of Sciences of the United States of America* **111**, 8619-8624.

[6] Kar, K.; Kubilius, J.; Schmidt, K.; Issa, E. B. and DiCarlo, J. J. (2019). Evidence that recurrent circuits are critical to the ventral stream's execution of core object recognition behavior. *Nature neuroscience* **22**, 974-983.
